# Probiotic acoustic biosensors for noninvasive imaging of gut inflammation

**DOI:** 10.1101/2024.09.23.614598

**Authors:** Marjorie T. Buss, Lian Zhu, Jamie H. Kwon, Jeffrey J. Tabor, Mikhail G. Shapiro

**Affiliations:** Division of Chemistry and Chemical Engineering, California Institute of Technology, Pasadena, CA, USA; Division of Biology and Biological Engineering, California Institute of Technology, Pasadena, CA, USA; Andrew and Peggy Cherng Department of Medical Engineering, California Institute of Technology, Pasadena, CA, USA; Howard Hughes Medical Institute, Pasadena, CA, USA; Ph.D. Program in Systems, Synthetic, and Physical Biology, Rice University, Houston, TX, USA; Department of Biosciences, Rice University, Houston, TX, USA; Department of Bioengineering, Rice University, Houston, TX, USA

## Abstract

Inflammatory bowel diseases (IBD) affect millions of people globally, result in severe symptoms, and are difficult to diagnose and monitor – often necessitating the use of invasive and costly methods such as colonoscopies or endoscopies. Engineered gut bacteria offer a promising alternative due to their ability to persist in the gastrointestinal (GI) tract and sense and respond to specific environmental signals. However, probiotics that have previously been engineered to report on inflammatory and other disease biomarkers in the Gl tract rely on fluorescent or bioluminescent reporters, whose signals cannot be resolved in situ due to the poor penetration of light in tissue. To overcome this limitation, we introduce probiotic biosensors that can be imaged in situ using ultrasound – a widely available, inexpensive imaging modality providing sub-mm spatial resolution deep inside the body. These biosensors are based on the clinically approved probiotic bacterium *E. coli* Nissle, which we engineered to transiently colonize the GI tract, sense inflammatory biomarkers, and respond by expressing air-filled sound-scattering protein nanostructures called gas vesicles. After optimizing biomolecular signaling circuits to respond sensitively to the biomarkers thiosulfate and tetrathionate and produce strong and stable ultrasound contrast, we validated our living biosensors in vivo by noninvasively imaging antibiotic-induced inflammation in mice. By connecting cell-based diagnostic agents to ultrasound, this “diagnostic yogurt” will make it easier, cheaper, and less painful to diagnose and monitor IBD or other GI conditions.

## INTRODUCTION

Inflammatory bowel diseases (IBD) affect millions of people^1,2^, result in severe life-altering symptoms^3,4^, and are caused by complex genetic and environmental factors leading to chronic intestinal inflammation^5,6^. Current methods to diagnose and monitor IBD, such as colonoscopies and biopsies, can be invasive, unpleasant, and costly^7^, causing delays in diagnosis and treatment that can lead to complications and reduce the effectiveness of therapeutic interventions^8–14^. Other less invasive methods such as blood and stool tests often lack specificity due to inflammation elsewhere in the body and the short-lived nature of many disease-associated intestinal molecules. Moreover, they do not provide spatial information on the location of inflammation within the intestines, which can influence treatment decisions^15–17^. A method to noninvasively visualize inflammatory biomarkers in situ in the gastrointestinal (GI) tract could greatly facilitate the monitoring, understanding, and diagnosis of IBD and other GI diseases^18–20^.

Engineered probiotic bacteria could help diagnose IBD by colonizing the GI tract and sensing and reporting on specific signals in their environment^21–23^. Indeed, engineered bacteria have been developed to monitor intestinal biomarkers such as thiosulfate^24^, tetrathionate^24,25^, nitric oxide^26,27^, calprotectin^28^, and bleeding^29^. However, most of these probiotics report their findings through fluorescent or colorimetric reporters in the feces, which lack information about where in gut the inflammation takes place and require specialized laboratory equipment and protocols (e.g. flow cytometry) that may be difficult to translate to the clinic. Bioluminescent reporter strains^28,30^ can be imaged in situ in mice, but have limited resolution due to light scattering and do not scale to larger animals or humans due limited light penetration^31^. Bioluminescent bacteria have been coupled with a pill-based wireless electronic device^29^, but this approach is complex, uses a large pill (∼4.5 cm) that would not be easy to swallow by humans, and provides limited spatiotemporal information due to its rapid passage through the GI tract.

As an alternative readout, ultrasound readily propagates through tissue, allowing images to be acquired at several-centimeter depth with 100 micron-level spatial resolution^31^. Moreover, ultrasound imaging is inexpensive and ubiquitously available^32^, increasing the translatability and potential impact of diagnostics that use ultrasound as a readout. Recently, it was demonstrated that bacteria can generate ultrasound contrast by expressing acoustic reporter genes (ARGs), which result in the production of air-filled protein nanostructures called gas vesicles that scatter sound waves^33,34^. However, ARG expression has not yet been incorporated into biosensors to detect specific molecular signals or used to visualize GI-colonizing bacteria.

We hypothesized that we could use ARG-expressing probiotic cells as the basis for a “diagnostic yogurt” that could be ingested by a patient, transiently populate the GI tract, sense an inflammatory biomarker, and produce acoustic contrast that could be detected with a simple, noninvasive ultrasound scan (**Fig. 1**). To test this concept, we engineered the clinically approved probiotic *Escherichia coli* Nissle 1917 (EcN) to express ARGs in response to small molecule biomarkers of GI inflammation – focusing on thiosulfate and tetrathionate^24^ – and developed protocols for their ultrasound detection in vivo after GI colonization. We optimized our acoustic biosensors through multiple rounds of genetic engineering so that they would produce detectable ultrasound contrast under physiologically relevant conditions (e.g. 37ºC, low biomarker concentrations). Finally, we validated mouse models of antibiotic-induced inflammation that exhibit elevated thiosulfate, suggesting this may be a general IBD biomarker, and noninvasively imaged inflammation in these mice using our optimized acoustic thiosulfate sensor.

**Figure 1:**
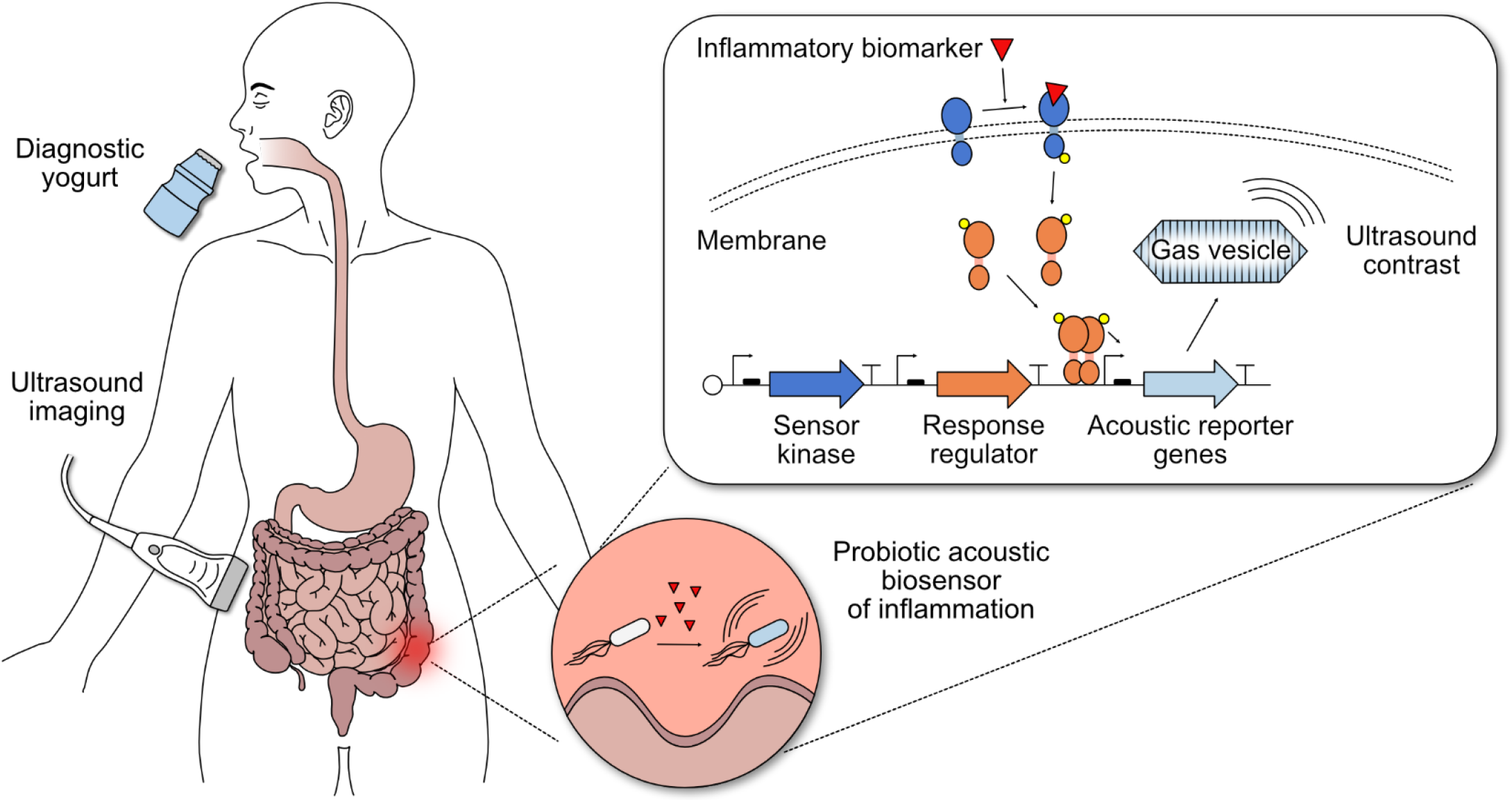
Concept of probiotic biosensors for ultrasound imaging of gastrointestinal (GI) inflammation. Patients consume yogurt containing diagnostic probiotic bacteria that transiently populate the GI tract. Using two-component systems (TCSs), these engineered bacteria sense inflammatory biomarkers, such as thiosulfate and tetrathionate, and express acoustic reporter genes (ARGs) that encode for gas vesicles (GVs) in response. Binding of the biomarker to the membrane sensor kinase protein triggers transcription of ARGs from a specific promoter via phosphorylation of a cytoplasmic response regulator protein; in the absence of the biomarker, phosphatase activity of the membrane sensor kinase keeps the system repressed. ARG-expressing bacteria can be detected via ultrasound imaging in patients, enabling ultrasound imaging of GI inflammation.

## RESULTS

### Engineered biosensing pathways enable the production of ultrasound contrast in response to inflammatory biomarkers

Thiosulfate (S_2_O_3_^2-^) and tetrathionate (S_4_O_6_^2-^) are biomarkers for intestinal inflammation arising from altered gut sulfur metabolism during colitis. Thiosulfate is generated when host cells detoxify hydrogen sulfide (H_2_S)^35^, whose production is thought to be upregulated during intestinal inflammation^36–39^. Reactive oxygen species present during inflammation further convert thiosulfate into the transient product tetrathionate^40^. Thiosulfate and tetrathionate have been shown to be elevated in DSS-induced^24^ and *Salmonella*-induced^25^ mouse models of colitis, respectively.

To develop living biosensors of thiosulfate and tetrathionate, we started with previously described fluorescent sensors of these molecules based on two-component systems (TCS) from *Shewanella*^24^. We aimed to replace the GFP in these constructs with ARGs to enable ultrasound imaging of the sensor bacteria in situ rather than detection of GFP in the feces via flow cytometry. We used a recently optimized ARG construct called bARG_Ser_, derived from *Serratia sp*^34^, placing it downstream of the promoter driven by *thsR* or *ttrR* response regulators, which are activated by thiosulfate or tetrathionate through *thsS* or *ttrS* sensor kinase receptors, respectively. We implemented these genetic pathways on a single medium-copy plasmid, with the TCS components constitutively expressed and with the Axe-Txe stability cassette^41^ to form the plasmids thsSR-bARG_Ser_ and ttrSR-bARG_Ser_ (**Fig. 2a**).

**Figure 2:**
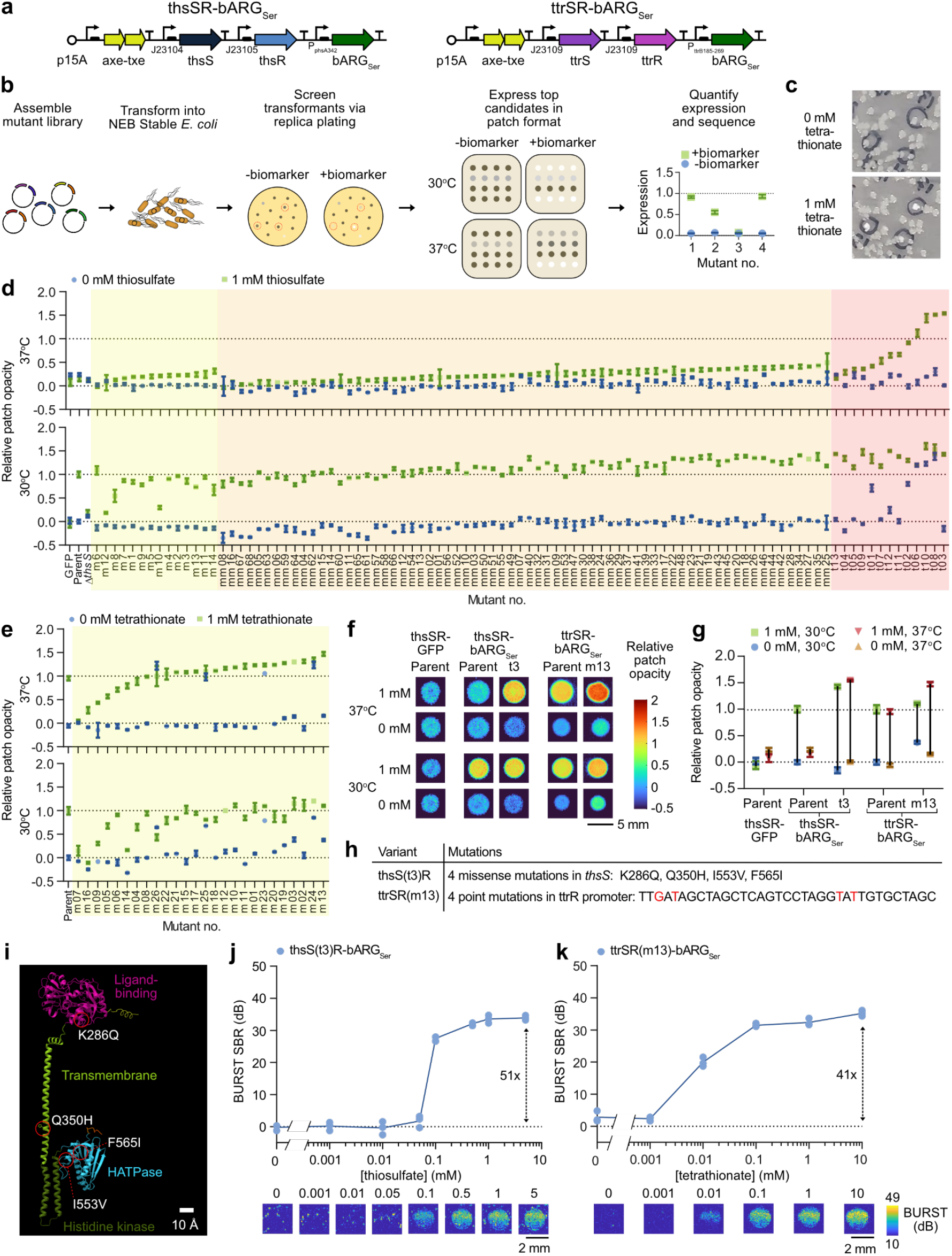
Optimization of thiosulfate and tetrathionate sensors with ARGs as the output. (**a**) Plasmid diagrams of thiosulfate (thsSR-bARG_Ser_) and tetrathionate (ttrSR-bARG_Ser_) sensors with ARGs from *Serratia* sp. ATCC 39006 (bARG_Ser_) as the output.(**b**) Protocol for screening for optimized variants of these sensors. A mutant library is assembled via error-prone PCR or PCR with semi-random primers followed by Gibson assembly and is then transformed into NEB Stable *E. coli*. Transformants are screened by replica plating onto plates with 0 or 1 mM thiosulfate or tetrathionate. Colonies that are more opaque, corresponding to higher ARG expression, at 1 mM than at 0 mM are further characterized in patch format at 0 and 1 mM thiosulfate/tetrathionate at 30°C and 37°C. Patch opacity is quantified and plotted to compare variants. (**c**) Representative photographs of replica plates containing colonies at 0 and 1 mM tetrathionate, where circled colonies were more opaque at 1 mM than at 0 mM. (**d**) Relative patch opacities from screening thsSR-bARG_Ser_ variants mutated in the response regulator promoter (m01-m14, yellow shading), in both the response regulator and the sensor kinase promoters (mm01-mm68, orange shading), and in the sensor kinase gene *thsS* (t01-t13, red shading). (**e**) Relative patch opacities from screening ttrSR-bARG_Ser_ variants mutated in the response regulator promoter (m01-m26, yellow shading). (**f-g**) Representative images (f) and quantification in terms of relative opacity (g) of patches of the best-performing variants compared to the parent construct and a GFP control in NEB Stable *E. coli*. In (d-g), a relative patch opacity of 0 corresponds to the parent construct at 0 mM thiosulfate/tetrathionate and a relative patch opacity of 1 corresponds to the parent construct at 1 mM thiosulfate/tetrathionate at 30°C. Variants were ordered from lowest to highest relative patch opacity at 37°C within each screening group. (**h**) Summary of mutations in the best-performing variants; see **Table S1** for full sequences. (**i**) AlphaFold 3^Ref. 89^ structural prediction of *thsS(t3)* colored by predicted domains with the four missense mutations indicated by red circles; see **Fig. S3** for more views of structural predictions. (**j-k**) Representative BURST ultrasound images (bottom) and quantification of the signal-to-background ratio (SBR) (top) of the best variants for thsSR-bARG_Ser_ (j) and ttrSR-bARG_Ser_ (k) at varying thiosulfate/tetrathionate concentrations in *E. coli* Nissle (EcN) at 37°C in liquid culture. See **Fig. S4** for corresponding xAM data. Cells were cast in agarose phantoms at 10^9^ cells/mL for ultrasound imaging. In (d, e, g), points represent the average of 2-4 biological replicates and error bars represent the standard deviation. In (j-k), lines represent the mean of 3 biological replicates which are each averaged over 2 technical replicates

Using these constructs, we observed ARG expression in *E. coli* in response to thiosulfate and tetrathionate, but the initial ultrasound signal was lower than with an arabinose-inducible positive control construct pBAD-bARG_Ser_ (**Fig. S1**). To improve performance, we engineered our genetic constructs using rapid mutagenesis and screening, relying on the optical opacity of gas vesicle-expressing bacterial colonies and replica-plating onto thiosulfate- or tetrathionate-containing media (**Fig. 2b-c**). In our screens, colonies that were more opaque with 1mM thiosulfate/tetrathionate than without were picked and characterized in patch format at 30°C and 37°C. First, we tuned the expression levels of response regulators *thsR*/*ttrR* by mutagenizing their constitutive promoters using semi-random primers (**Fig. S2a**). This yielded several variants of thsSR-bARG_Ser_ with slight improvements in opacity fold-change, but all variants still performed poorly at 37°C compared to 30°C (**Fig. 2d**). For ttrSR-bARG_Ser_, this approach yielded several variants with higher opacity at 37°C and 1 mM tetrathionate – in particular the m13 variant (**Fig. 2e**). Similar trends were observed when mutating the GFP-expressing versions of the sensors (**Fig. S2b-f**).

Mutating the promoters driving both *thsR* and *thsS* simultaneously led to improved thiosulfate-sensing performance at 30°C and 37°C, but performance remained poor at 37°C, leading us to hypothesize that the membrane sensor kinase *thsS* might not express well at the higher temperature. To address this issue, we mutated *thsS* using error-prone PCR and found several variants with high opacity at 37°C and 1 mM thiosulfate. In particular, the *thsS(t3)* variant exhibited low opacity at 0 mM and very high opacity at 1 mM thiosulfate at both 37°C and 30°C (**Fig. 2d**). Overall, the thsS(t3)R-bARG_Ser_ variant, which contained 4 missense mutations (K286Q, Q350H, I553V, and F565I) and 2 silent mutations in *thsS*, exhibited 10.5 times higher opacity at 37°C and 1 mM thiosulfate than its parent construct (**Fig. 2f-h, Fig. S3, Table S1**). The mutated residues were distributed across the ligand-binding, transmembrane, and catalytic domains of the protein (**Fig. 2i**). The ttrSR(m13)-bARG_Ser_ variant, which contained 4 point mutations in the *ttrR* constitutive promoter, had 1.5 times higher opacity at 37°C and 1 mM tetrathionate than its parent (**Fig. 2f-h, Table S1**).

We characterized the ultrasound contrast produced by our two improved biosensors in EcN cultured at 37ºC with a range of thiosulfate and tetrathionate concentrations, observing increasing signal in response to increasing biomarker concentrations (**Fig. 2j-k, Fig. S4**). Using BURST imaging, a highly sensitive and specific ultrasound pulse sequence for gas vesicle detection^42^, thsS(t3)R-bARG_Ser_ and ttrSR(m13)-bARG_Ser_ exhibited maximal fold-changes of 51 and 41, respectively, in response to their cognate analytes. This switching performance is high compared to many inducible biosensors in the literature^24,27,28,43^. Similar trends were observed under xAM imaging, a complementary, non-destructive ultrasound imaging mode also popular for gas vesicle detection^44^ (**Fig. S4**).

### Recombinase-based switching increases sensor signal in response to biomarkers

To increase ultrasound signal and biomarker sensitivity even further, we incorporated a recombinase into our optimized two-component sensors. We placed the serine integrase Bxb1 downstream of our *thsR*-activated promoter, and placed bARG_Ser_ downstream of the strong constitutive promoter P7^45^, which we flanked with Bxb1 recognition sequences *attB* and *attP*. We included a temperature-responsive terminator upstream of Bxb1 to reduce leaky expression outside the body below 37°C^46^ (**Fig. 3a**). Before cells are exposed to thiosulfate, the P7 promoter points towards two strong terminators and bARG_Ser_ is not expressed. When cells are exposed to thiosulfate, the thsS(t3)R pathway triggers Bxb1 expression, which catalyzes irreversible site-specific recombination between *attB* and *attP*, flipping the direction of P7 and resulting in constitutive bARG_Ser_ expression. We named this construct thsS(t3)R-Bxb1_P7-bARG_Ser_

**Figure 3:**
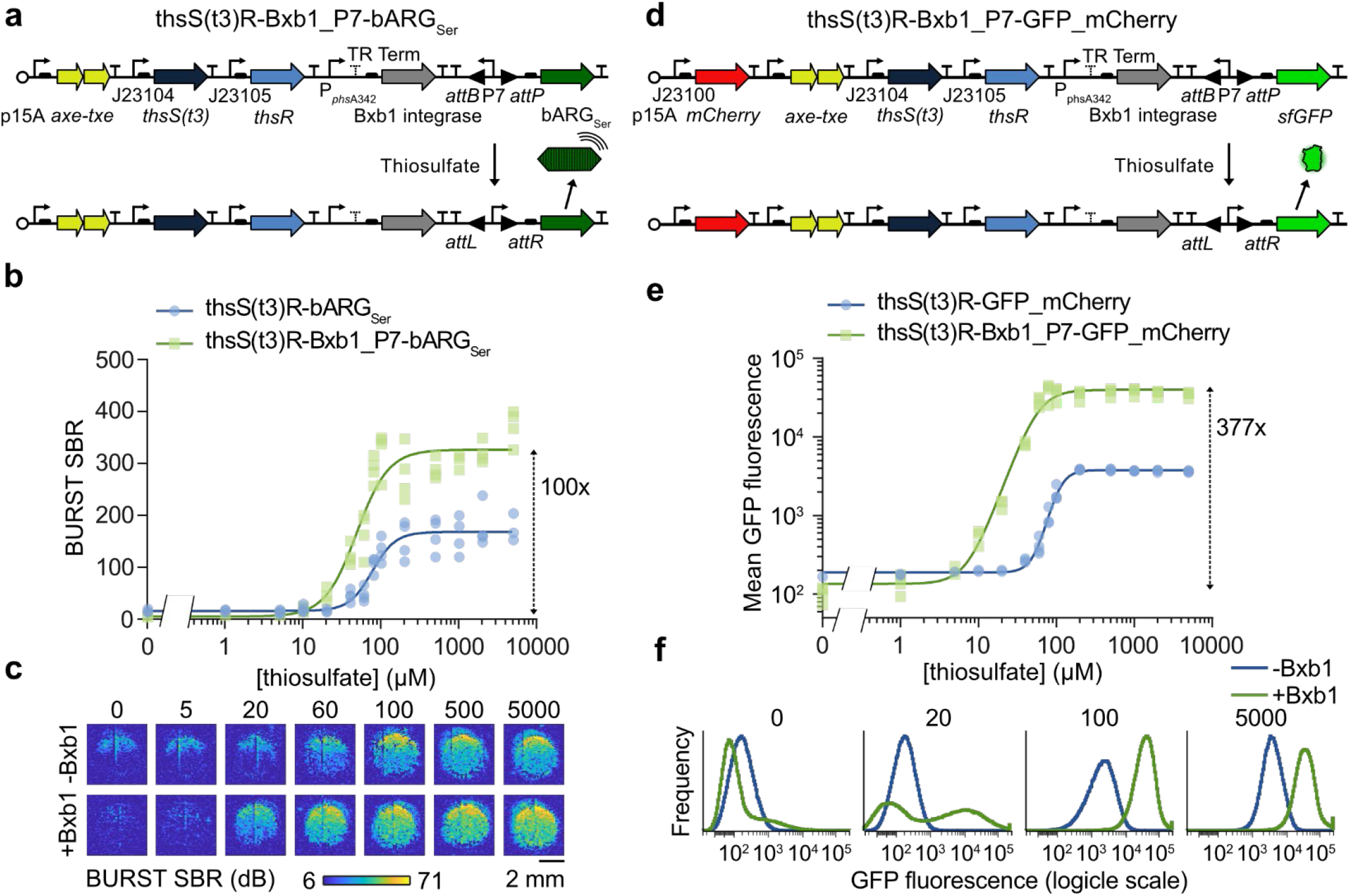
Increasing sensor activation with addition of an integrase-based switch. (**a**) Plasmid diagram of the optimized thiosulfate sensor thsS(t3)R-bARG_Ser_ with an integrase-based switch to create thsS(t3)R-Bxb1_P7-bARG_Ser_. Addition of thiosulfate induces expression of the Bxb1 integrase which causes irreversible site-specific recombination between the *attP* and *attB* sites that reverses the orientation of the strong constitutive P7 promoter to activate bARG_Ser_ expression. (**b-c**) BURST signal-to-background ratio (SBR) (b) and representative images (c) of the optimized thiosulfate sensor with and without the Bxb1 integrase-based switch at varying thiosulfate concentrations. See **Fig. S6a-b** for the corresponding xAM data. (**d**) Plasmid diagram of the optimized thiosulfate sensor thsS(t3)R-GFP_mCherry with an integrase-based switch to create thsS(t3)R-Bxb1_P7-GFP_mCherry. (**e-f**) Mean GFP fluorescence measured via flow cytometry (e) and representative histograms (f) of the optimized thiosulfate sensor with and without the Bxb1 integrase-based switch at varying thiosulfate concentrations. In (b) and (e), points represent biological replicates (N=4), curves represent fits to the Hill equation (see **Table S2** for fitted parameters), and numbers next to dashed arrows indicate maximal fold changes. All strains were grown on plates with varying concentrations of thiosulfate at 37°C (see **Fig. S5** for images of the plates) and suspended in PBS for ultrasound imaging and flow cytometry; for ultrasound imaging, cells were cast in agarose phantoms at a concentration of 5 × 10^8^ cells/mL. See **Fig. S6c-j** for the corresponding data for the tetrathionate sensor.

We compared thsS(t3)R-Bxb1_P7-bARG_Ser_ and thsS(t3)R-bARG_Ser_ in EcN at 37°C, measuring the BURST and xAM ultrasound signals after exposing the cells to a range of thiosulfate concentrations (**Fig. 3b-c, Fig. S5a, Fig. S6a-b**). The recombinase-based biosensors produced 2.5-fold higher ultrasound signal in saturating (100 µM) thiosulfate compared to non-recombinase constructs, resulting in a maximal fold-change of 100. The recombinase-based circuits also showed higher thiosulfate sensitivity, shifting the half-maximum Hill equation constant for BURST signal from 76.4 to 48.2 µM (**Table S2**).

We also implemented the Bxb1 switching approach in the fluorescent protein version of the thiosulfate sensor, forming plasmid thsS(t3)R-Bxb1_P7-GFP_mCherry (**Fig. 3d**). Flow cytometry of thsS(t3)R-Bxb1_P7-GFP_mCherry versus thsS(t3)R-GFP_mCherry EcN after inducing with a range of thiosulfate concentrations at 37°C demonstrated increased GFP fluorescence in response to thiosulfate (19.2-fold higher at 100 µM thiosulfate), increased sensitivity (the Hill constant decreased from 105 to 57.2 µM), and distinct negative and positive (“unflipped” and “flipped”) GFP populations with addition of the Bxb1 switch (**Fig. 3e-f, Fig. S5b, Table S2**).

We obtained similar results when implementing the Bxb1 switch in the ARG and fluorescent versions of our tetrathionate sensor (**Fig. S5c-d, Fig. S6c-j, Table S2**). These results demonstrate that addition of a Bxb1 switch with the strong constitutive promoter P7 improves the performance of two-component system EcN biosensors with both acoustic and fluorescent outputs.

### Probiotic biosensors colonize the GI tract and respond to their cognate biomarkers

To prepare for disease biosensing in vivo, we first aimed to demonstrate that EcN-based probiotic agents could colonize the GI tract and produce ultrasound contrast in response to externally supplied small molecules. To reduce colonization resistance by native fauna, we treated the mice with streptomycin for two days prior to probiotic administration^47–50^ and used EcN with a spontaneous mutation in the *rpsL* gene conferring streptomycin resistance (**Table S3**). Initially, we orally gavaged arabinose-inducible pBAD-bARG_Ser_ EcN (**Fig. S7**) into mice receiving arabinose in their drinking water (**Fig. 4a**). We used fecal colony plating to confirm stable GI colonization for at least 3 days (**Fig. 4b**), a duration sufficient for the probiotics to sense their environment and produce gas vesicles (high levels of which are reached within 24 hours of induction)^34^. We imaged the mice using a home-built ultrasound scanning system that enables rapid large-scale imaging of the abdomen (**Fig. S8a**), and BURST* ultrasound – a version of the BURST^42^ pulse sequence that we optimized for GI probiotic imaging (as described in Methods). Subsequently, we sacrificed the animals for confirmatory ex vivo imaging of their intestines (**Fig. S8b**).

**Figure 4:**
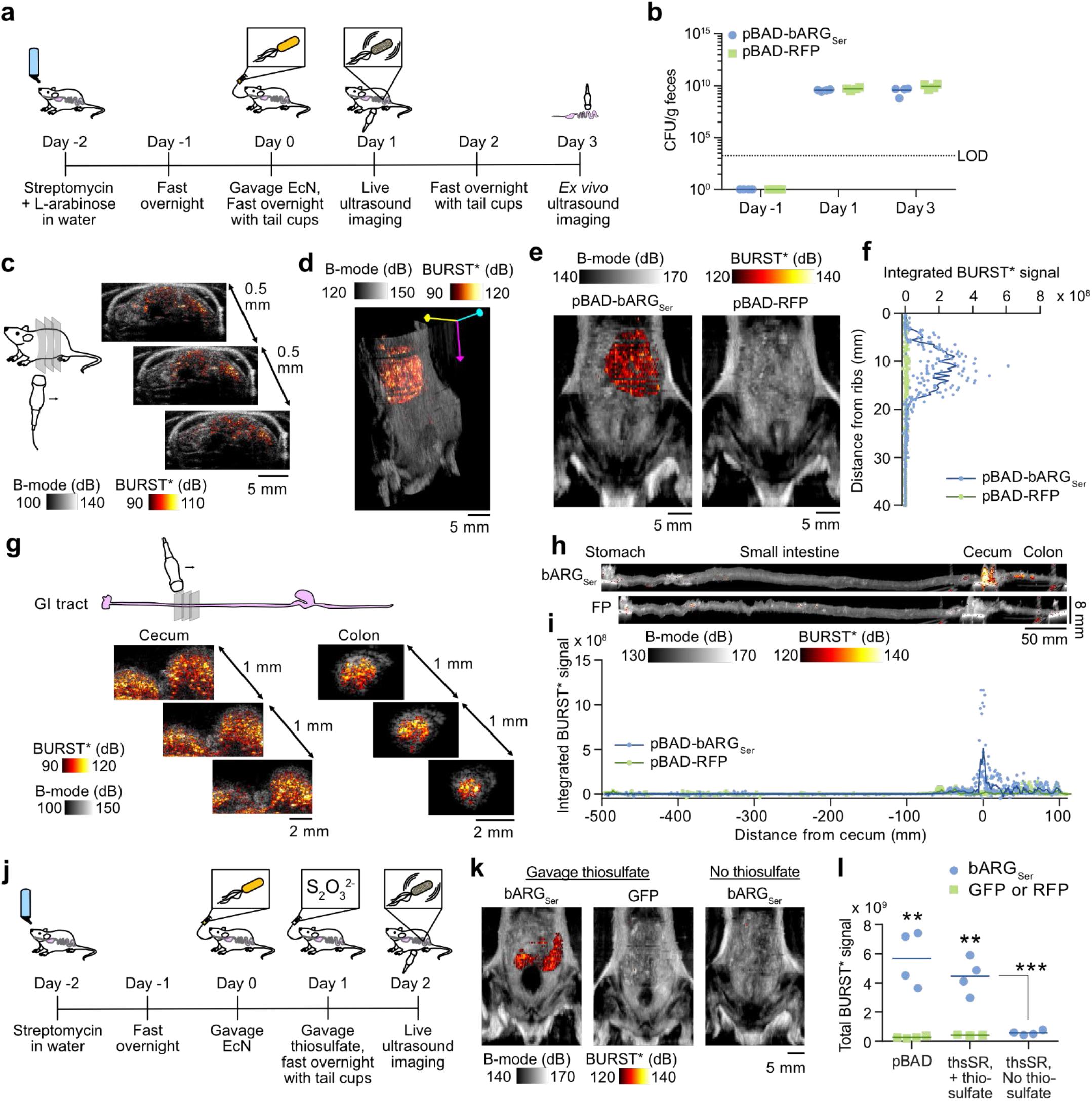
Imaging ARG expression by EcN in the GI tract in response to L-arabinose and thiosulfate. (**a**) Experimental design for testing L-arabinose-inducible bARG_Ser_ expression in EcN in vivo. Mice were given water containing L-arabinose and streptomycin for 2 days, and the streptomycin-resistant EcN containing pBAD-bARG_Ser_ or the control plasmid pBAD-RFP were orally administered. After fasting with tail cups overnight, mice were scanned with ultrasound. After allowing another day for re-expression, mice were sacrificed and their intestines were scanned with ultrasound ex vivo. (**b**) Colony forming units (CFU) per gram of feces before, 1 day after, and 3 days after administration of the L-arabinose-sensing EcN. Limit of detection (LOD) was 1.7 × 10^3^ CFU/g feces. (**c**) Representative 2-D images from scanning a mouse colonized by pBAD-bARG_Ser_ EcN where the BURST* image (hot scale) was thresholded and overlaid onto the B-mode image (greyscale). Images were acquired as transverse planes spaced by 0.5 mm from the rib cage to the tail; see **Fig. S8a** for the imaging setup. (**d**) 3-D projection of the BURST* (hot scale) and B-mode (greyscale) images acquired from scanning a representative mouse colonized by pBAD-bARG_Ser_ EcN. See **Supplementary Videos 1-2** for more views of the 3-D images. (**e**) Representative ultrasound images overlaying the integrated BURST* signal over the depth onto the integrated B-mode signal over the depth one day after administration of the L-arabinose-sensing EcN. See **Fig. S9a** for images of all mice. (**f**) Quantification of the integrated BURST* signal versus the distance from the ribs one day after administration of the L-arabinose-sensing EcN. (**g**) Representative 2-D ex vivo images from scanning the GI tract of a mouse colonized by pBAD-bARG_Ser_ EcN where the BURST* images (hot scale) were thresholded and overlaid onto the B-mode images (greyscale) for the colon (left) and cecum (right). Images were acquired as transverse planes spaced by 1 mm from the stomach to the rectum; see **Fig. S8b** for the imaging setup. (**h**) Representative ex vivo ultrasound images of intestines 3 days after administration of the L-arabinose-sensing EcN. The integrated BURST* signal over the width was overlaid onto the integrated B-mode signal over the width. Note that the pixels are not isotropic due to the large length to diameter ratio of the intestines (see scale bars). See **Fig. S9b** for images of all intestines. (**i**) Quantification of the integrated BURST* signal versus the length of the intestines relative to the cecum. (**j**) Experimental design for testing thiosulfate-sensing EcN strains in vivo. Mice were treated with streptomycin for 2 days, EcN strains were orally administered, and the next day 1 M thiosulfate was orally administered or omitted from control mice. After allowing a day for bARG_Ser_ expression, mice were imaged with ultrasound. (**k**) Representative ultrasound images overlaying the integrated BURST* signal onto the integrated B-mode signal over the depth for thiosulfate-sensing EcN one day after oral gavage of thiosulfate, or without any external thiosulfate for control mice. See **Fig. S11** for images of all mice and colonization data. (**n**) Total BURST* ultrasound signal imaged in mice colonized by L-arabinose-sensing (pBAD) or thiosulfate-sensing (thsSR) EcN strains with either bARG_Ser_ or GFP as the output. Asterisks represent statistical significance by two-tailed, unpaired Student’s t-tests (** = p < 0.01, *** = p < 0.001). p-values from left to right: 0.00120628, 0.0026301, 0.00078501. For (b), (f), (i), and (l), each point represents a biological replicate (N=4 for all except N=3 for thsSR-GFP in (l)) and lines represent the mean.

Scans of mice acquired one day after gavage of pBAD-bARG_Ser_ EcN revealed high levels of ultrasound contrast in the cecum, which was not present in control mice administered with the fluorescent pBAD-RFP EcN control (**Fig. 4c-f, Fig. S9a, Supplementary Videos 1-2**). Ex vivo imaging three days after gavage confirmed that the BURST* signal was concentrated primarily in the cecum and colon (**Fig. 4g-i** and **Fig. S9b**), where resistant EcN are known to colonize in streptomycin-treated mice^30^. Negligible mutational silencing was observed in EcN plated from the feces during the course of this experiment (**Fig. S9c**). BURST* signal was also detected directly in feces by ultrasound imaging (**Fig. S10**).

We next aimed to confirm that our thiosulfate-sensing acoustic biosensor EcN cells could be detected via ultrasound in mice directly fed with thiosulfate. After 2 days of streptomycin treatment, the thsS(t3)R-bARG_Ser_ and thsSR-GFP probiotics were orally administered to populate the GI tract (**Fig. 4j, S11a-b**). One day after gavage of the biosensors, 1 M thiosulfate was orally administered, or omitted from control mice, and the next day all mice were imaged with ultrasound. In the presence of thiosulfate, mice colonized with the probiotic acoustic biosensor exhibited significantly higher BURST* signal in the cecum compared to mice that did not receive the corresponding biomarker or were colonized by control fluorescent strains (**Fig. 4k-l**). The amount of contrast scaled with the amount of administered thiosulfate (**Fig. S11c-e**). These results demonstrated that our probiotic EcN cells are capable of populating the mouse GI tract in vivo and functioning as an acoustic biosensor of thiosulfate when this analyte is present at sufficient concentrations.

### Streptomycin-treated mice exhibit dysbiosis-associated inflammation

To test our biosensors in a disease context, we set out to establish and validate a mouse model of inflammatory dysbiosis. Since streptomycin treatment can cause inflammation^51^, we aimed to determine whether mice treated with this antibiotic exhibit elevated intestinal thiosulfate levels. Accordingly, we treated mice with streptomycin sulfate or the control antibiotic chloramphenicol for 5 days (**Fig. 5a**). The streptomycin-treated mice exhibited signs of dysbiosis and inflammation, including looser stools and greater weight loss compared to chloramphenicol-treated controls (on top of acute weight reduction caused by fasting with tail cups between days 2 and 3) (**Fig. 5b**). Using ion chromatography-mass spectrometry (IC-MS) to quantify thiosulfate (**Fig. S12**), we found that fecal thiosulfate levels were elevated in streptomycin-treated mice compared to chloramphenicol-treated (**Fig. 5c**) or untreated (**Fig. S13a-b**) animals. In separate experiments, we verified that sulfate on its own, at the same or higher level as present in our streptomycin sulfate solution, did not affect fecal thiosulfate (**Fig. S14**), consistent with literature^52^. These thiosulfate levels were higher than those measured in mice treated with dextran sulfate sodium (DSS), another agent known to cause GI inflammation^53^ (**Fig. S13a-b**).

**Figure 5:**
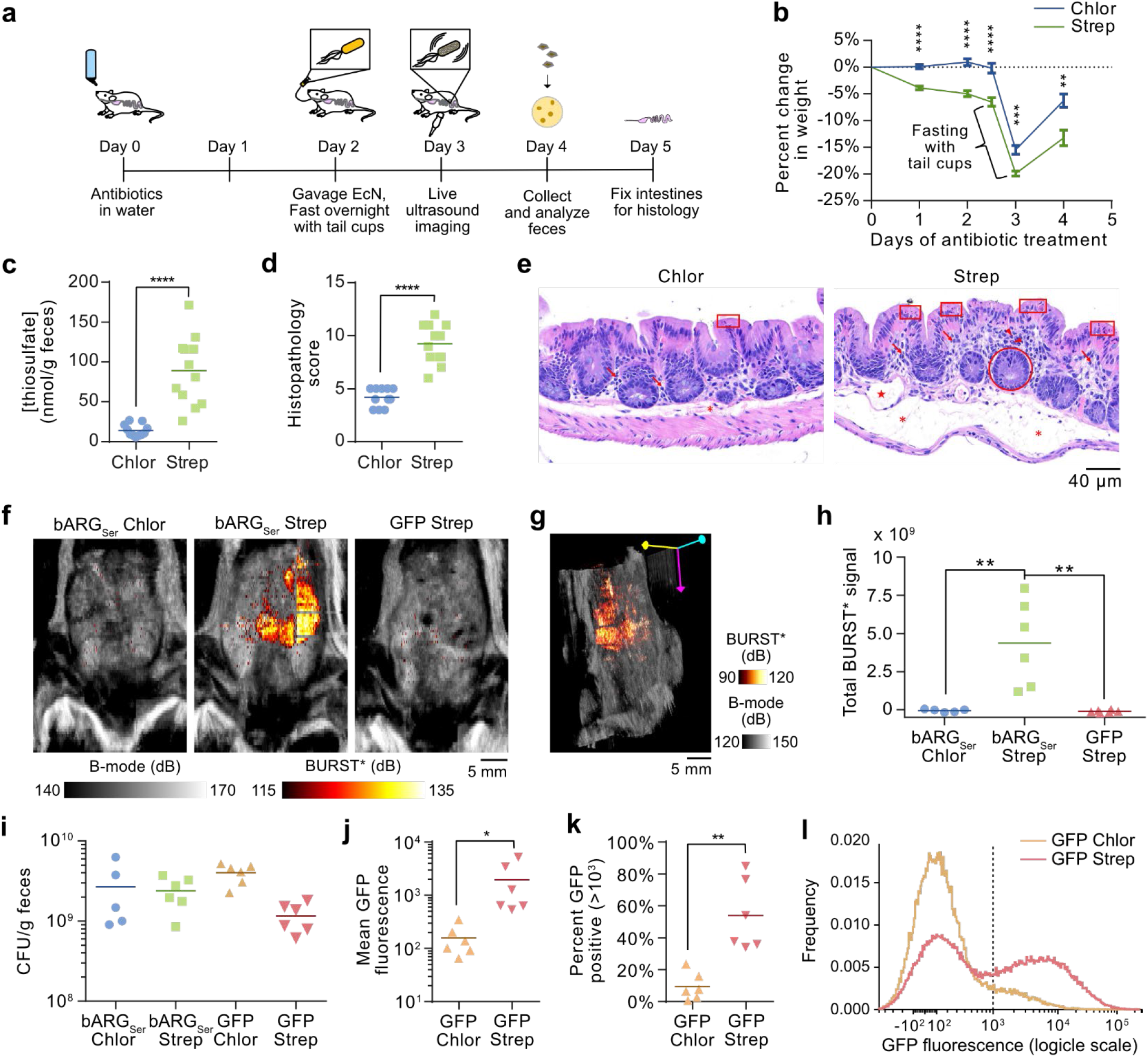
Ultrasound imaging of thiosulfate sensor activation during antibiotic-induced inflammation. (**a**) Experimental design for testing EcN strains containing plasmids for the optimized integrase-based switch thiosulfate sensors, thsS(t3)R-Bxb1_P7-bARG_Ser_ or thsS(t3)R-Bxb1_P7-GFP_mCherry. (**b**) Percent change in weight after addition of chloramphenicol or streptomycin to the water of the mice. P-values from left to right: 1.309050e-07, 7.544644e-07, 3.185578e-05, 0.000135693, 0.00171773. N = 11 for chloramphenicol-treated mice and N = 12 for streptomycin-treated mice. (**c**) Thiosulfate concentration in nmol per gram feces measured via IC-MS 4 days after antibiotic administration. See **Fig. S12** for IC-MS chromatograms and standard curves. P-value: 1.17999e-05. (**d-e**) Histopathology scores (d) and representative images (e) of H&E-stained sections of cecal tissue from mice sacrificed on day 5 after antibiotic administration. Abnormalities are indicated in red: mucosal epithelial cell death and degeneration (box), mucosal crypt hyperplasia (circle), mucosal/submucosal edema (asterisk), mononuclear infiltrates (arrow), granulocytic infiltrates (arrowhead), dilated lymphatic (star). See **Fig. S15** for the scoring by category and additional images. P-value: 5.38198E-08 (k). (**f**) Representative ultrasound images overlaying the integrated BURST* signal onto the integrated B-mode signal for mice treated with chloramphenicol and colonized with thsS(t3)R-Bxb1_P7-bARG_Ser_ EcN (bARG_Ser_ Chlor), for mice treated with streptomycin and colonized with thsS(t3)R-Bxb1_P7-bARG_Ser_ EcN (bARG_Ser_ Strep), and for mice treated with streptomycin and colonized with thsS(t3)R-Bxb1_P7-GFP_mCherry EcN (GFP Strep) on day 3. See **Fig. S16** for all images of mice on day 3. (**g**) Representative 3-D projection of BURST* and B-mode images of a mouse treated with streptomycin and colonized with thsS(t3)R-Bxb1_P7-bARG_Ser_ EcN. See **Supplementary Videos 3-5** for more views of 3-D projections from this experiment. (**h**) Total BURST* signal in all mice in the groups from (c) on day 3. P-values from left to right: 0.0068, 0.0029. (**i**) Colony forming units (CFU) per gram of feces on day 4 for all mice in the groups from (f), plus mice treated with chloramphenicol and colonized with thsS(t3)R-Bxb1_P7-GFP_mCherry EcN (GFP Chlor). (**j-l**) Mean GFP fluorescence, percent GFP positive events (> 10^3^), and aggregate histograms of GFP fluorescence from flow cytometry analysis of feces from mice colonized with thsS(t3)R-Bxb1_P7-GFP_mCherry EcN and treated with chloramphenicol (GFP Chlor) or streptomycin (GFP Strep) on day 4. P-values: 0.0466 (g), 0.0010 (h). Asterisks represent statistical significance by two-tailed, unpaired Student’s t-tests (* = p < 0.05, ** = p < 0.01, *** = p < 0.001, **** = p < 0.0001). Points represent biological replicates (N=5 for bARG_Ser_ Chlor, N=6 for bARG_Ser_ Strep, N=6 for GFP Chlor, and N=6 for GFP Strep), lines represent means, and error bars represent the standard error of the mean.

Blinded histological analysis of the cecum tissue from mice sacrificed after 5 days of streptomycin treatment revealed significantly more inflammatory pathology, including mononuclear infiltrates, granulocytic infiltrates, and mucosal crypt hyperplasia, compared to controls (**Fig. 5d-e, Fig. S15**). These results confirm that streptomycin treatment leads to intestinal dysbiosis^54^ and inflammation^51^ and elevates fecal thiosulfate. Quantitatively, the concentration of thiosulfate measured in the feces (∼90 µM in streptomycin-treated mice, assuming an approximate density of 1 g feces per mL^55^) and initial flow cytometry measurements (**Fig. S13c-e**) suggested that our recombinase-based biosensors would be the most suitable agents for detecting this inflammatory process with ultrasound.

### Probiotic acoustic biosensors enable noninvasive imaging of GI inflammation in a mouse model of inflammatory dysbiosis

To test whether our optimized thiosulfate biosensors can visualize streptomycin-induced inflammation, we orally administered the thsS(t3)R-Bxb1_P7-bARG_Ser_ and thsS(t3)R-Bxb1_P7-GFP_mCherry EcN probiotics two days after the start of dysbiosis induction (**Fig. 5a**). Ultrasound imaging on day 3 revealed strong BURST* signal in the cecum of streptomycin-treated mice that received the acoustic biosensor; this signal was not present in chloramphenicol-treated controls that received the same biosensor (which carries chloramphenicol resistance), nor in streptomycin-treated mice that received the fluorescent version of the sensor (**Fig. 5f-h, Fig. S16, and 3D renderings in Supplementary Videos 3-5**). Plating of feces on day 4 confirmed that both biosensor strains colonized streptomycin-and chloramphenicol-treated mice well (**Fig. 5i**). Corroborating our acoustic biosensor results, flow cytometry of the feces obtained on day 4 from mice administered with the fluorescent biosensor showed increased GFP fluorescence and a larger GFP-positive population in streptomycin-treated animals (**Fig. 5j-l**).

The observed BURST* signal in individual animals was positively correlated with the measured weight loss and thiosulfate levels, which were themselves positively correlated with the histopathology score (**Fig. S17**). These results demonstrate the in-situ activation of thiosulfate-sensing acoustic biosensors in a disease model, allowing a specific inflammatory biomarker to be imaged noninvasively with ultrasound. To show that our biosensing approach can be generalized beyond streptomycin, we also developed a model of inflammatory dysbiosis caused by piperacillin and obtained similar ultrasound imaging results (**Fig. S18, Table S3**).

## DISCUSSION

In this study, we introduced, optimized and validated the first probiotic biosensors that can be used to visualize GI inflammation noninvasively with ultrasound. We showed that these biosensors could be imaged 1-3 days after oral administration and responded robustly to multiple compounds (arabinose, tetrathionate, thiosulfate) to produce ultrasound contrast robustly detectable above the backdrop of intestinal tissue and luminal contents. We demonstrated that a probiotic acoustic biosensor of thiosulfate – optimized through several rounds of mutagenesis, screening, and circuit design – could be used to noninvasively visualize inflammatory dysbiosis caused by two different antibiotic treatments, with in vivo imaging data correlating with disease severity.

Because ultrasound imaging systems are relatively inexpensive and widely available^32^, we anticipate that the development of ARG-producing probiotic biosensors will help facilitate the clinical adoption of “diagnostic yogurts”. Compared with current methods to diagnose and monitor IBD, such as colonoscopies and stool tests, our approach provides an alternative that is simultaneously noninvasive and in situ, bridging the gap between molecular-level information and anatomical imaging. Our approach is also highly modular, as ARGs can be coupled with virtually any transcriptional biosensor, enabling the noninvasive imaging of a wide range of disease biomarkers or other molecular species important to health. For instance, probiotic bacteria could be engineered to sense and noninvasively report on a range of important species in the GI tract, including bile acids^56,57^, formate^58,59^, lactate^60^, and short-chain fatty acids^61^, to better understand their role in health and disease or diagnose and track human conditions.

Specific to the biosensing targets of this study, our experiments emphasize the role of antibiotics in causing dysbiosis and intestinal inflammation through host-microbial interactions^54,62–67^. Streptomycin, commonly used to assist *E. coli* colonization in rodent studies^48,49^, induces mild inflammation^51^ and dysbiosis^68–71^ that increases intestinal thiosulfate levels, likely due to a bloom of *Enterobacteriaceae* and other facultative anaerobes which metabolize sulfomucins and sulfur-containing amino acids, producing sulfide as a byproduct that is detoxified by the host to thiosulfate^36–38,72–75^. Chloramphenicol-treated mice might represent mice with a healthier GI tract by suppressing the growth of SRB^76,77^ and other bacteria that contribute to sulfide generation. Overall, our results suggest that streptomycin treatment is a mouse model for dysbiosis-associated intestinal inflammation, with thiosulfate as a biomarker for disease severity.

Outside of inflammatory biosensing, our results suggest that ARGs could be useful for the noninvasive study and monitoring of a larger variety of commensal microbes, pathogens, or engineered bacteria in the GI tract^78^. ARGs could be expressed in different species of bacteria to visualize their locations and gene expression over time in response to different perturbations to the GI tract. For example, to better understand the role of gut microbes in gastrointestinal cancers^79^, colonization of different bacteria in tumors along the GI tract could be monitored using ARGs. Additionally, the spatiotemporal expression of virulence genes by pathogenic bacteria^80^ or commensal colonization factors by commensal bacteria^81^ could be noninvasively monitored by linking them with ARG expression to better understand their roles in causing GI infections or aiding colonization.

We envision several improvements to our technology. First, given the role of antibiotics in causing dysbiosis and inflammation, the colonization of sensor bacteria should not be dependent on antibiotic treatment. To instead obtain high levels of stable colonization of sensor bacteria, the bacteria could be engineered to metabolize an external carbon source that cannot be used by members of the native microbiome; this strategy has been successfully used in *Bacteroides* by engineering it to consume the polysaccharide porphyran^82^. Second, the detection of bARG_Ser_ expression in the GI tract relies on collapse of gas vesicles through BURST imaging due to the high background and non-uniformity from intestinal contents and tissue, limiting imaging sessions to at most once per day to allow time for gas vesicle re-expression. To overcome this limitation, bARG_Ser_ could be engineered to produce more nonlinear contrast that can be detected above background in the GI tract so that it can be imaged nondestructively with amplitude modulation pulse sequences^44,83^. Additionally, to complement thiosulfate and tetrathionate, additional sensors could be engineered to sense human-validated biomarkers such as calprotectin^28^ and tested in models of inflammation that display more localized patterns of disease, such as TNBS-induced colitis^84^. Finally, to facilitate the clinical adoption of this technology, parallel advances in ultrasound scanning technology could enable volumetric imaging^85,86^ or be adapted for continuous long-term disease monitoring by using adhesive ultrasound devices^87,88^. With these improvements, diagnostic yogurts could become a valuable new flavor of biotechnology.

## METHODS

### Molecular biology

All plasmids were assembled using reagents from New England Biolabs (NEB) for Gibson Assembly and from fragments generated via PCR with Q5 High Fidelity DNA polymerase (NEB). Assemblies were transformed into NEB Stable *E. coli* via electroporation for plasmid preparation and maintenance, and all plasmids were verified with commercial Sanger sequencing (Laragen) or whole-plasmid sequencing (Primordium Labs). Plasmids containing the thiosulfate and tetrathionate TCS components, pKD236-4b (encoding *thsS*), pKD237-3a-2 (encoding *thsR*), pKD238-1a (encoding *ttrS*), and pKD239-1g-2 (encoding *ttrR*), were gifts from Jeffrey Tabor (Rice University). Components for the recombinase-based switch, including Bxb1, P7, attB, and attP, were amplified from plasmids from Abedi et al^46^. The plasmid backbone containing a chloramphenicol resistance gene and the p15A origin of replication, as well as arabinose-inducible components (araC and pBAD promoter), bARG_Ser_, and Axe-Txe were amplified from plasmids from Hurt et al^34^. The pOSIP plasmid kit used for clonetegration was a gift from Drew Endy and Keith Shearwin (Addgene kit # 1000000035). Genomic modifications to *E. coli* Nissle (EcN) were verified by colony PCR with OneTaq DNA polymerase (NEB), gel purification of bands, and sequencing (Laragen or Primordium Labs). Integrated DNA Technologies synthesized other genes and all PCR primers.

### Materials and Media

LB media was prepared using 10 g/L Bacto tryptone (BD Biosciences), 5 g/L Bacto yeast extract (BD Biosciences), and 5 g/L NaCl; LB-agar plates were prepared with the addition of 15 g/L Bacto agar (BD Biosciences). M9 media was prepared using 1x M9 salts (Sigma-Aldrich), 0.4% v/v glycerol, 0.2% w/v Bacto casamino acids (BD Biosciences), 2 mM MgSO_4_, and 0.1 mM CaCl_2_; M9-agar plates were prepared with the addition of 15 g/L Bacto agar (BD Biosciences). Sodium thiosulfate pentahydrate (Sigma-Aldrich) and potassium tetrathionate (Sigma-Aldrich) solutions were freshly prepared in water and used within a day. DSS (36,000 - 50,000 MW, colitis grade, MP Biomedicals) solutions were prepared fresh every 2 days. Chloramphenicol (Sigma-Aldrich), streptomycin sulfate (Sigma-Aldrich), and piperacillin sodium salt (Cayman Chemical Company) were prepared as 1000X stock solutions and stored at -20°C for in vitro experiments or were prepared fresh for animal experiments. All other chemicals were of analytical grade and commercially available.

### Mutant library generation and screening

For site-directed mutagenesis of constitutive Anderson promoters, semi-random primers (**Fig. S2a**) were used for PCR with Q5 High Fidelity DNA polymerase (NEB). For error-prone PCR of *thsS*, Taq DNA polymerase (NEB) with the standard reaction buffer (NEB), 0.2 mM dATP/dGTP, 1 mM dCTP/dTTP, 0.5 µM primers, 5.5 mM MgCl_2_, and 0.2 mM or 0.5 mM MnCl_2_ was used. After assembly via Gibson Assembly, mutant libraries were transformed via electroporation into NEB Stable *E. coli*. Various 10-fold dilutions of the outgrowth were plated onto LB-agar plates with antibiotics (30 µg/mL chloramphenicol + 100 µg/mL streptomycin) to ensure evenly spaced colonies for replica plating. Transformant colonies were transferred to M9-agar plates with antibiotics with and without the appropriate inducer (1 mM thiosulfate or tetrathionate) via replica plating, and replica plates were grown at 30°C or 37°C for 20-24 hours. Colonies that were more opaque or that had higher GFP expression at 1 mM than at 0 mM inducer were picked from the plate with 0 mM inducer into LB media with antibiotics and grown at 37°C overnight. One µL of the overnight culture was then dropped onto M9-agar plates with varying thiosulfate or tetrathionate concentrations to make patches, and patch plates were grown at 30°C or 37°C for 20-24 hours. ARG-expressing patches were quantified in terms of opacity by acquiring white trans images with the standard filter using a Bio-Rad ChemiDoc MP gel imager. GFP-expressing patches or colonies were quantified by acquiring an image using blue epi illumination and a 530/28 filter. For quantifying patch opacity, ROIs were drawn around patches and the background directly adjacent to the patch using ImageJ. The mean pixel intensity of the background ROI next to the patch was subtracted from the mean pixel intensity of the patch ROI to correct for nonuniformity in the background of the white trans image. The background-subtracted mean pixel intensities were then normalized so that a value of 0 represented the opacity of the parent strain at 0 mM inducer and 30°C, and a value of 1 represented the opacity of the parent strain at 1 mM inducer and 30°C. These values were termed “relative patch opacity.”

Because *E. coli* Nissle (EcN) appeared to mutate plasmids during transformation, all EcN transformants had to be screened. Sequence-verified plasmids purified from NEB Stable *E. coli* cultures were transformed into EcN via electroporation. The resulting transformant colonies were screened by streaking onto plates with antibiotics (30 µg/mL chloramphenicol, 100 µg/mL streptomycin, and/or 25 µg/mL piperacillin) and with and without the appropriate inducer (1 mM thiosulfate or tetrathionate). Only colonies which expressed GFP/bARG_Ser_ on the inducer plate were picked from the plate without inducer into LB media with antibiotics. LB cultures were grown at 37°C or 30°C overnight, and cryostocks were prepared by gently mixing 500 µL of the overnight culture with 500 µL of 30% glycerol and placing at -80°C.

### In vitro characterizations

To characterize strains in vitro, the appropriate cryostock was used to inoculate LB + 0.4% glucose + antibiotics (30 µg/mL chloramphenicol, 100 µg/mL streptomycin, and/or 25 µg/mL piperacillin) which was grown overnight at 37°C and 250 rpm. For characterizations in liquid culture, the overnight culture was used to inoculate 25-50 mL of M9 media + antibiotics in 250 mL baffled flasks at an initial OD_600_ of 0.05. Once the OD_600_ of the M9 culture reached 0.1-0.3, 1 mL aliquots were distributed in 15 mL tubes and induced with thiosulfate, tetrathionate, or L-arabinose or left uninduced. The 1 mL cultures were grown at 37°C and 250 rpm for 20-24 hours before being placed at 4°C or on ice until analyzed by ultrasound imaging or flow cytometry. For characterizations on solid media, 1 µL from the overnight culture was dropped onto M9 plates + antibiotics and the desired inducer concentrations. Plates were grown at 37°C for 20-24 hours and the resulting patches were scraped off using inoculating loops into PBS and kept on ice or at 4°C until ultrasound imaging or flow cytometry.

### Ultrasound imaging of in vitro bacterial samples and feces

To prepare phantoms of bacterial cells for ultrasound imaging, wells were cast with a custom 3D-printed mold using 1% (w/v) agarose in PBS, which was degassed by incubating at 65-75°C for at least 16 hours. The culture or cell suspension to be imaged was diluted to 2x the final desired cell concentration in a volume of 100 µL in PBS on ice. The 100 µL of cells in PBS was placed on a heating block at 42°C for approximately one minute, 100 µL of molten 1% agarose in PBS at 42°C was gently mixed in, and the mixture was pipetted into the wells of the phantom in duplicate, taking care to avoid bubbles. To prepare phantoms of feces for ultrasound imaging, 10 µL of molten 1% agarose in PBS at 55°C was pipetted into a well and a fecal pellet was quickly pushed into the well into the molten agarose. More molten agarose was pipetted on top of the fecal pellet to completely fill the well and any air pockets.

Once solidified, the phantoms were submerged in PBS, and ultrasound images were acquired using a Verasonics Vantage programmable ultrasound scanning system and an L22-14v (**Fig. 2, S1, S4, S7, S10**) or an L22-14vX (**Fig. 3, S6**) 128-element linear array transducer. The L22-14v transducer had a center frequency of 18.5 MHz with 67%-6-dB bandwidth and an elevation focus of 6 mm. The L22-14vX transducer had a center frequency of 18.0 MHz with >60%-6-dB bandwidth and an elevation focus of 8 mm. Both transducers had an element pitch of 100 µm and an elevation aperture of 1.5 mm. The transducer was attached to a custom-made manual translation stage to move between samples.

B-mode and xAM images were acquired using the same parameters as described previously^34^: the frequency and transmit focus were set to 15.625 MHz and 5 mm, respectively, and each image was an average of 50 accumulations. B-mode imaging was performed with a conventional 128-ray-lines protocol, where each ray line was a single pulse transmitted with an aperture of 40 elements. xAM imaging was performed using a custom sequence detailed previously^44^ with an angle of 19.5° and an aperture of 65 elements. With the L22-14v transducer, BURST images were acquired as a series of pAM^83^ images where the focus was set to 6 mm and the frequency was set to 18.0 MHz with 2 waveform cycles (3 half-cycles plus a half-cycle equalization), and the voltage was set to 1.6V for the first 10 frames and 25V for the last 46 frames. With the L22-14vX transducer, BURST images were acquired as a series of B-mode images, where three 32-aperature focused beams were acquired at a time to improve the frame rate by a factor of 3, and where the first image was acquired at 1.6 V and the last 7 images were acquired at 20V. The focus was set to 6 mm and the frequency was set to 18.0 MHz with 3 half-cycles and a half-cycle equalization.

For image processing and analysis, custom beamforming scripts were applied on-line to reconstruct images from the acquired RF data. BURST images were calculated as the first collapsing frame minus the last collapsing frame (**Fig. S19, Supplementary Note 1**). Circular ROIs were drawn around the sample wells and around the background regions in the phantom without wells in MATLAB. The signal-to-background ratio was calculated as the mean pixel intensity of the sample ROI divided by the mean pixel intensity of the background ROI. Conversion to decibels (dB) was calculated as 20*log10(SBR). For display, images were normalized by dividing by the average background signal of all images being compared and setting the lower and upper limits of the colormaps to be the same, where the lower limit was equal to a constant A and the upper limit was equal to a constant B times the maximum pixel intensity divided by the average background out of all images being compared; images were then converted to dB. For BURST images, A=3 and B=1 with the L22-14v transducer, and A=2 and B=0.5 with the L22-14vX transducer. For xAM images, A=2 and B=0.5 with the L22-14v transducer, and A=2 and B=0.4 with the L22-14vX transducer.

### Animal procedures

All animal experiments were approved by the California Institute of Technology Institutional Animal Care and Use Committee (IACUC). Animals were housed in a facility maintained at 71-75°F and 30-70% humidity, with a lighting cycle of 13 hours on & 11 hours off (light cycle 6:00 AM - 7:00 PM). All mice were 6-10 week-old female Balb/c (**Fig. 4, S9, S10, S11**) or male C57BL/6 (**Fig. 5, S13, S14, S15, S16, S17, S18**) mice obtained from Jackson Labs. For antibiotic, DSS, and inducer treatments administered in the drinking water *ad libitum*, the appropriate compounds were dissolved in water and filtered with a 0.2 µm membrane; the water was freshly prepared at least every 2 days. Standard rodent diet (5053 - PicoLab Rodent Diet 20) was provided *ad libitum* unless mice were being fasted. Oral gavage was performed using a 20 gauge 1.5” length animal feeding needle and with a volume of 200 uL. Prior to gavage, mice were fasted in individual housing for 2-3 hours. EcN strains were prepared for gavage by growing an overnight culture in LB + 0.4% glucose + antibiotics at 37°C from a cryostock, diluting the overnight culture 1:100 into 50 mL of fresh M9 or LB + 0.4% glucose + antibiotics in 250 mL baffled flasks, incubating at 37°C and 250 rpm until the OD_600_ reached 0.4-0.6, pelleting the culture at 3500g at room temperature for 10 min, and suspending in an appropriate volume of sterile PBS for an OD_600_ of 20. Feces were collected by placing mice individually in empty cages without food or bedding for approximately 20-30 minutes.

### In vivo ultrasound imaging

Prior to imaging, mice were fasted overnight with tail cups in individual housing with access to water only to reduce the amount of material and background ultrasound signal in the GI tract^89,90^. Shortly before imaging, mice were anesthetized with 2% isoflurane, the tail cups were removed, and the abdominal fur was removed with shaving and Nair. Approximately 5 minutes before imaging, 1.2 µg atropine in a volume of 60 µL saline was injected subcutaneously on each side of the abdomen to slow movement of intestines due to peristalsis^91,92^. Mice were placed prone and partially submerged in water onto a thin film of mylar (2.5 µm thickness, Chemplex, catalogue number 100) which was acoustically transparent. A nose cone provided isoflurane and kept the head out of the water, white a heating lamp was used to regulate the body temperature. The ultrasound transducer (L22-14v or L22-14vX) was submerged using a probe cover (Protek, part number 1-519-2450) and attached to a BiSlide computer-controlled 3-D translatable stage (Velmex). The transducer was positioned 6 mm below the mylar at an angle of 15° from the vertical to minimize specular reflection, and at a distance from the mylar to center the 6 mm focus on the intestines. See **Fig. S8a** for a diagram.

A custom MATLAB script was used to control the ultrasound system and the 3-D translatable stage at the same time so that the entire abdominal area of the mouse was automatically scanned. Starting at the rib cage, the transducer was moved in three 9.6-mm steps across the body and in eighty 0.5-mm steps lengthwise down the body to the tail. At each spot, BURST imaging was performed, giving a total of 240 BURST acquisitions per mouse which took around 15 min to acquire. BURST acquisitions were a series of B-mode images, where three 32-aperature focused beams were acquired at a time to improve the frame rate by a factor of 3^Ref. 42^, with the first image taken at 1.6 V and the last seven images taken at the collapsing voltage. The focus was set to 6 mm and the frequency was set to 18.0 MHz. With the L22-14v transducer (used for in vivo data in **Fig. 4, S9, and S11**), 3 transmit waveform cycles were used and the collapsing voltage was 25V. With the L22-14vX transducer (used for in vivo data in **Fig. 5, S16, and S18**), 2 transmit waveform cycles were used and the collapsing voltage was 20V due to its higher pressure-to-voltage curve.

BURST* images were calculated by subtracting the first collapsing frame from the second because the ARG-specific signal was characterized by an increase in signal from the first to the second collapsing frame and using more frames was confounded by tissue motion (**Fig. S19, Supplementary Note 1**). Acquisitions during which breathing occurred were manually excluded. The first collapsing frame was used for the B-mode images. At each transverse plane, the three acquisitions across the body were stitched together to form one B-mode image and ROIs were drawn around the GI tract in each transverse plane B-mode image. 3-D BURST* and 3-D B-mode images were formed by stitching all 240 acquisitions together. For 3-D BURST* images, pixels were set to zero anywhere breathing occurred or outside of the ROIs. For display in 3-D, 3-D BURST* and 3-D B-mode arrays were converted to dB and loaded into napari^93^ where the BURST* image rendering and blending was set to “additive.” For display in 2-D, the B-mode signal was summed over the depth of the mouse from 4 to 8 mm, and the BURST* signal was summed over the depth of the entire ROI. The 2-D integrated BURST* image was thresholded, converted to dB, and overlaid onto the dB-converted 2-D integrated B-mode image. To calculate the total BURST* signal, all pixel intensities in the 3-D BURST* array inside the ROI excluding breathing were summed.

### Ex vivo ultrasound imaging

Prior to imaging, mice were fasted overnight with tail cups in individual housing with access to water only to reduce the amount of material and background ultrasound signal in the GI tract. Mice were euthanized by sedating with isoflurane and performing cervical dislocation, and the entire GI tract (stomach to rectum) was quickly removed and linearized by removing mesenteric tissue, taking care not to rip the intestines. The intestines were submerged in a degassed water bath where the stomach, cecum, and rectum were pinned down on platforms so that the entire GI tract was in approximately a straight line at constant depth. The platforms consisted of acoustic absorbers (Precision acoustics) on top and magnets embedded into the platforms on the bottom so that the platforms would stay submerged and could be moved to stretch out the intestines (whose length varied between mice) using another external magnet on the bottom of the water bath. The L22-14v ultrasound transducer was attached to a BiSlide computer-controlled 3-D translatable stage (Velmex) and was positioned 8 mm above the intestines at an angle of 15° from the vertical to minimize specular reflection. See **Fig. S8b** for a diagram.

A custom MATLAB script was used to control the ultrasound system and the 3-D translatable stage at the same time so that the entire GI tract was automatically scanned. Starting at the stomach, the transducer was moved in approximately six hundred 1-mm steps lengthwise down to the rectum, which took around 30 min. At each spot, BURST acquisitions were obtained as a series of B-mode images, where three 32-aperature focused beams were acquired at a time to improve the frame rate by a factor of 3, with the first image taken at 1.6 V and the last seven images taken at 25V. The focus was set to 8 mm and the frequency was set to 18.0 MHz with 3 transmit waveform cycles.

As with the in vivo images, ex vivo BURST* images were calculated by subtracting the first collapsing frame from the second (**Fig. S19, Supplementary Note 1**), and the first collapsing frame was used for the B-mode images. For display in 2-D, both BURST* and B-mode images were summed over the width of the transducer (9.6 mm). The 2-D integrated BURST* image was thresholded, converted to dB, and overlaid onto the dB-converted 2-D integrated B-mode image. To calculate the integrated BURST* signal along the length of the GI tract, all BURST* pixel intensities were summed between a depth of 4 and 12 mm.

### Processing fecal samples for downstream analysis

Approximately 50-150 mg of feces were collected per mouse and were stored on ice immediately after collection. Feces were homogenized at 100 mg feces per mL in sterile ice-cold PBS using vortexing and an MP Biomedical FastPrep 24 Tissue Homogenizer set to 5 m/s for 20-40 seconds. One hundred µL of the homogenized feces were saved for plating, 100-200 µL were saved for flow cytometry, and the rest were centrifuged at 10,000g and 4°C for 20 min. The supernatant was filtered through a 0.2 µm cellulose acetate membrane and was frozen at -80°C until analysis via IC-MS. The 100-200 µL that were saved for flow cytometry were mixed 1:1 with PBS containing 2 mg/mL chloramphenicol, filtered through a 40 µm membrane (Falcon Cell Strainers), and incubated at 37°C and 250 rpm for one hour to allow fluorophore maturation while protein synthesis was inhibited by the 1 mg/mL chloramphenicol^24^. The mixture was then stored at 4°C until analysis via flow cytometry within 24 hours. The 100 µL that was saved for plating was diluted in 900 µL sterile PBS, and five more 10-fold serial dilutions were made and plated on agar plates with antibiotics (30 µg/mL chloramphenicol, 100 µg/mL streptomycin, and/or 25 µg/mL piperacillin) and with and without inducers (1 mM thiosulfate, 1 mM tetrathionate, or 0.1% L-arabinose) using the drop plate method^94^. Total colony counts were used to calculate colony forming units (CFUs) per gram of feces, and the number of colonies that failed to express reporter genes on plates with inducer was used to calculate the fraction mutant colonies.

### Flow cytometry

A MACSQuant VYB flow cytometer (Miltenyi Biotec) was used for all flow cytometry analysis with the following settings: low flow rate, medium mixing, 25-50 µL uptake volume, standard mode, chilled 96 rack, and a trigger by SSC with a threshold of 2.0. The Y2 channel was used for mCherry and the B1 channel was used for GFP. For analyzing in vitro samples, appropriate dilutions in PBS + 0.5% (w/v) BSA + 1 mg/mL chloramphenicol were prepared to target 10^6^ cells/mL. For analyzing fecal samples, the 50 mg/mL feces + 1 mg/mL chloramphenicol in PBS samples that were stored at 4°C were diluted to 2.5 mg/mL feces in PBS + 0.5% (w/v) BSA + 1 mg/mL chloramphenicol (1/20 dilution). Cytoflow^95^ was used with custom Python scripts for flow cytometry data analysis. Events were gated on FSC-A and SSC-A characteristic of *E. coli* and on positive mCherry fluorescence. The geometric mean was used to calculate mean GFP fluorescence, and the fraction of GFP positive cells was calculated as the number of events above 1000 in the B1 channel divided by the total number of gated events. For histograms, the number of bins was calculated according to the Freedman-Diaconis rule (with a minimum of 100 bins) and a logicle scale with parameters from Cytoflow was used for the x-axis.

### Ion chromatography mass spectrometry (IC-MS)

Thiosulfate was quantified via IC-MS using a Dionex Integrion HPIC system coupled to an ISQ EC Single Quadrupole Mass Spectrometer (Thermo Scientific). A Dionex AS-AP autosampler was used to inject 5 μL of sample into a 25 μL sample loop in push-partial mode. Ion chromatography was performed using a Dionex IonPac AS18 analytical column (2 × 250 mm) with a guard column (2 × 50 mm) at a flow rate of 0.25 mL/min. An eluent generator (with Dionex Cartridge EGC 500 KOH) was used to create the following KOH gradient: 12 to 44 mM from 0-5 min, 44 mM from 5-8 min, 44-52 mM from 8-10 min, 52 mM from 10-15 min, 52 to 12 mM from 15-15.05 min, and 12 mM from 15.05-22 min. A Dionex 2 mm AERS suppressor operated at 33 mA was used to remove the KOH before conductivity detection and negative ion mode mass spectrometry. The mass spectrometer was operated in component mode to scan for thiosulfate at m/z=113 using a source CID voltage of 10.

Thiosulfate standards were freshly prepared at concentrations ranging from 100 to 3000 nM in water with 1/10x PBS. The 100 mg/mL fecal filtrates in PBS to be analyzed with IC-MS were removed from -80°C, rapidly thawed in a room temperature water bath, and diluted 1/10 in water. Peaks were automatically integrated using Chromeleon 7.2.10 software with the Chromeleon 6 detection algorithm and custom parameters. Because all samples contained the same PBS concentration (1/10x), phosphate which was detected via conductivity was used as an internal standard; the thiosulfate peak area was normalized by the phosphate peak area to correct for variations in sampler injection volume.

### Histology of cecal tissue

Mice were euthanized by sedating with isoflurane and performing cervical dislocation. The GI tract was quickly removed and a portion of the cecum was excised, flushed with ice-cold PBS to remove the contents, and fixed in 10% neutral buffered formalin for 24 hours at 4°C. Fixed cecal tissues were transferred to 70% ethanol and stored at 4°C until being shipped to IDEXX Laboratories for paraffin-embedding, sectioning, and H&E staining. A pathologist performed blinded grading of microscopic changes twice independently as to severity using the International Harmonization of Nomenclature and Diagnostic (INHAND) Criteria grading system where 0 = no significant change, 1= minimal, 2 = mild, 3 = moderate, and 4 = severe^96^.

### Structural predictions

3D models for *thsS* and *thsS(t3)* were generated from their amino acid sequences using AlphaFold 3^Ref. 97^ using a seed value of 100. Structures were displayed and aligned using ChimeraX^98^.

### Data availability

Plasmids will be made available through Addgene upon publication. All other materials and data are available from the corresponding author upon reasonable request.

### Code availability

Ultrasound data acquisition and analysis code will be made available on the Shapiro Lab GitHub at https://github.com/shapiro-lab upon publication.

## ACKNOWLEDGEMENTS

The authors thank the Caltech Flow Cytometry & Cell Sorting Facility for assistance with flow cytometry, Dr. Nathan Dalleska & the Water and Environment Lab (WEL) at Caltech for assistance with IC-MS, and Dr. Genevieve Remmers & IDEXX Laboratories for assistance with histopathology sample preparation and interpretation. The authors would also like to thank Dr. Moshe Baruch for assistance with sending the initial thiosulfate and tetrathionate sensor plasmids. MTB was supported by an NSF GRFP fellowship. This research was supported by the Pew Charitable Trust and the Institute for Collaborative Biotechnologies (US Army, W911NF-19-D-0001 to M.G.S.). Related research in the Shapiro Laboratory is supported by the David and Lucille Packard Foundation and the Dreyfus Foundation. MGS is an investigator of the Howard Hughes Medical Institute.

## AUTHOR CONTRIBUTIONS

MTB and MGS conceived the study. MTB, LZ, and JHK planned and performed experiments. MTB analyzed data. JJT provided reagents and input on the research design and manuscript. MTB and MGS wrote the manuscript with input from all other authors. MGS supervised the research.

## COMPETING INTERESTS

The authors declare no competing financial interests.

## SUPPLEMENTARY INFORMATION

**Table S1:**
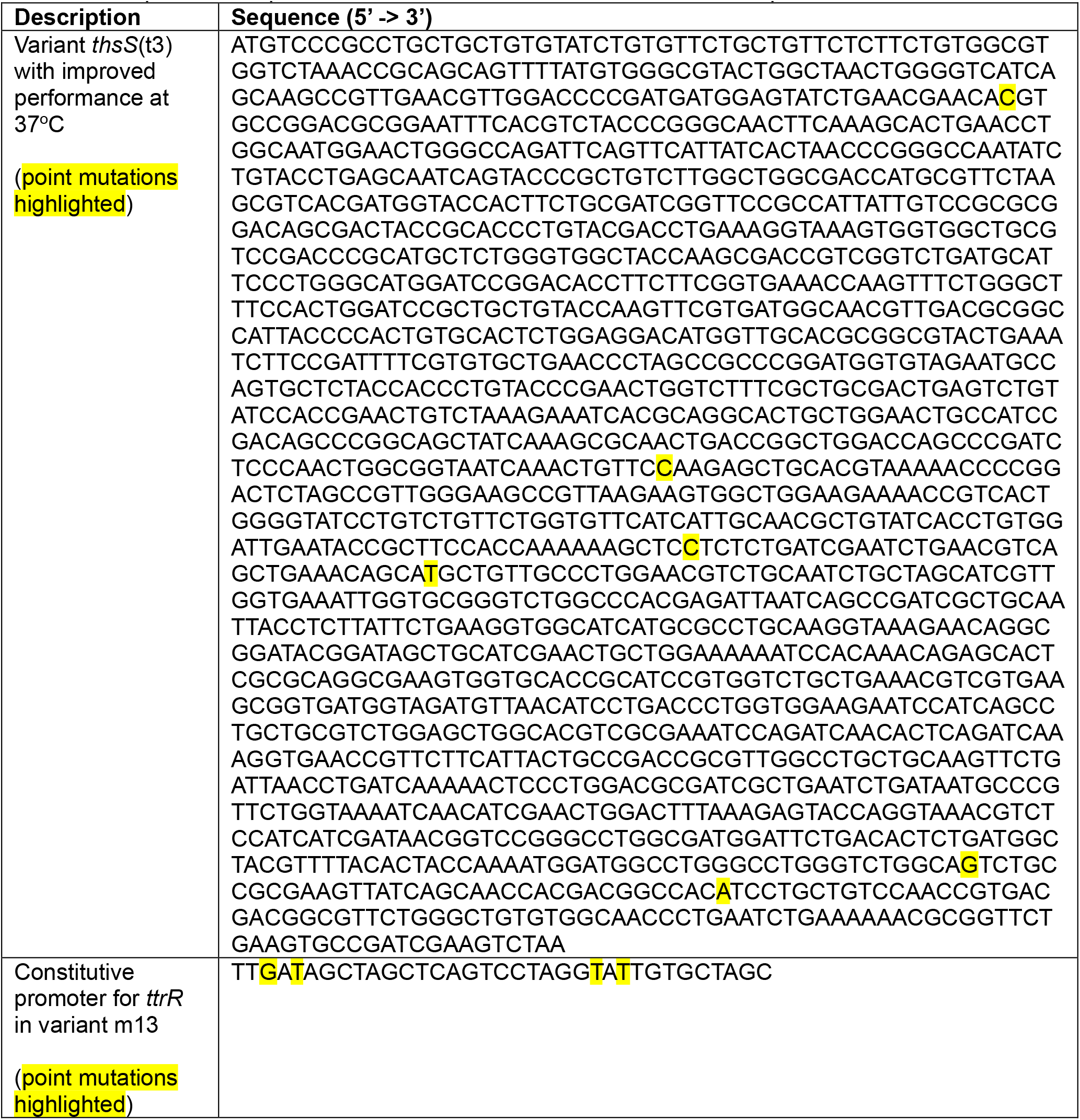
Sequences of optimized thiosulfate and tetrathionate sensor components.

**Table S2:** Parameters for fitting sensor characterization data from **Fig. 3** and **Fig. S6** to the Hill equation: 

| Output | BURST SBR |  | xAM SBR |  |
| --- | --- | --- | --- | --- |
| Strain | thsS(t3)R-bARG<sub>Ser</sub> | thsS(t3)R-Bxb1_P7-bARG<sub>Ser</sub> | thsS(t3)R-bARG<sub>Ser</sub> | thsS(t3)R-Bxb1_P7-bARG<sub>Ser</sub> |
| $A$ | 14.96 ± 0.9762 | 4.043 ± 0.3336 | 1.017 ± 0.01991 | 1.081 ± 0.03498 |
| $B$ | 152.1 ± 10.20 | 321.4 ± 16.83 | 5.416 ± 0.1221 | 12.88 ± 0.3666 |
| $K$ | 76.42 ± 6.386 | 48.23 ± 4.128 | 72.53 ± 2.080 | 54.93 ± 2.247 |
| $n$ | 2.912 ± 0.4397 | 2.126 ± 0.1304 | 3.002 ± 0.1860 | 2.654 ± 0.1585 |
| $R^2$ | 0.8769 | 0.9336 | 0.9834 | 0.9760 |

| Output | Mean GFP Fluorescence |  |
| --- | --- | --- |
| Strain | thsS(t3)R-GFP | thsS(t3)R-Bxb1_P7-GFP_mCherry |
| $A$ | 188.8 ± 5.021 | 135.3 ± 11.75 |
| $B$ | 3580 ± 104.9 | 39473 ± 2254 |
| $K$ | 105.0 ± 2.756 | 57.20 ± 3.942 |
| $n$ | 4.634 ± 0.2870 | 2.790 ± 0.1454 |
| $R^2$ | 0.9701 | 0.9180 |

| Output | BURST SBR |  | xAM SBR |  |
| --- | --- | --- | --- | --- |
| Strain | ttrSR(m13)-bARG<sub>Ser</sub> | ttrSR(m13)-Bxb1_P7-bARG<sub>Ser</sub> | ttrSR(m13)-bARG<sub>Ser</sub> | ttrSR(m13)-Bxb1_P7-bARG<sub>Ser</sub> |
| $A$ | 8.280 ± 0.5146 | 34.57 ± 1.830 | 1.034 ± 0.01942 | 2.124 ± 0.04621 |
| $B$ | 145.2 ± 10.37 | 203.7 ± 17.02 | 4.782 ± 0.1206 | 12.28 ± 0.4240 |
| $K$ | 32.77 ± 5.688 | 44.56 ± 7.114 | 25.58 ± 1.361 | 41.98 ± 2.996 |
| $n$ | 1.205 ± 0.1125 | 1.646 ± 0.2303 | 1.945 ± 0.1185 | 1.490 ± 0.08407 |
| $R^2$ | 0.9367 | 0.8859 | 0.9842 | 0.9801 |

| Output | Mean GFP Fluorescence |  |
| --- | --- | --- |
| Strain | ttrSR(m13)-GFP | ttrSR(m13)-Bxb1_P7-GFP_mCherry |
| $A$ | 57.80 ± 1.068 | 143.6 ± 5.553 |
| $B$ | 916.1 ± 18.95 | 31452 ± 1046 |
| $K$ | 34.53 ± 1.200 | 27.84 ± 1.271 |
| $n$ | 2.328 ± 0.08339 | 2.981 ± 0.1149 |
| $R^2$ | 0.9896 | 0.9668 |

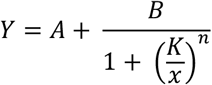

Here, *Y* is the sensor output (ultrasound signal or fluorescence) at the ligand (thiosulfate or tetrathionate) concentration *x* in µM. *A, B, K*, and *n* are constants where *A* is the minimum output with no ligand, *B* is the maximum output with saturating ligand concentration, *K* is ligand concentration that elicits a half-maximal response, and *n* is the Hill coefficient. Parameters are reported as the fitted value ± the standard error.

**Table S3:**
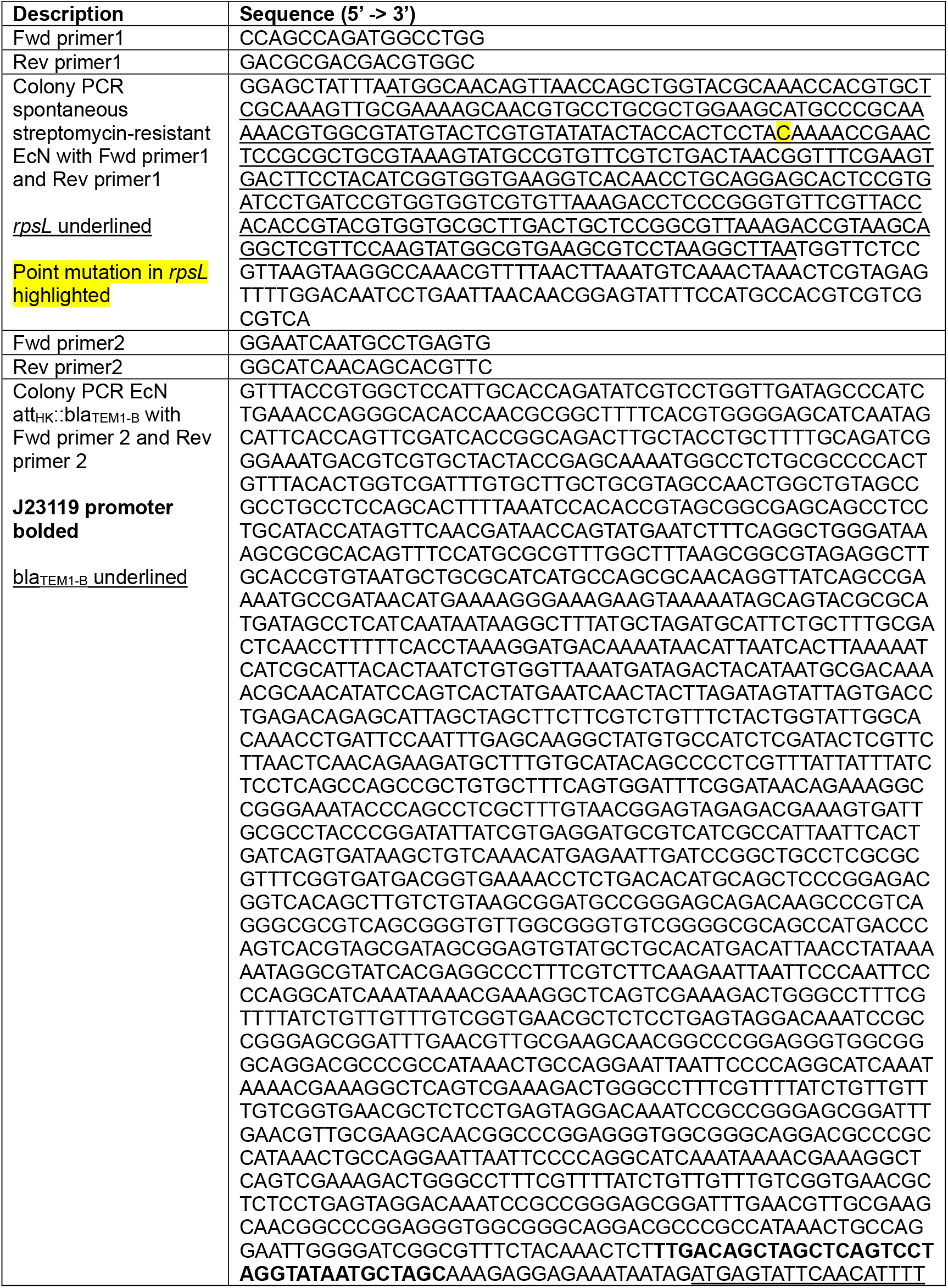
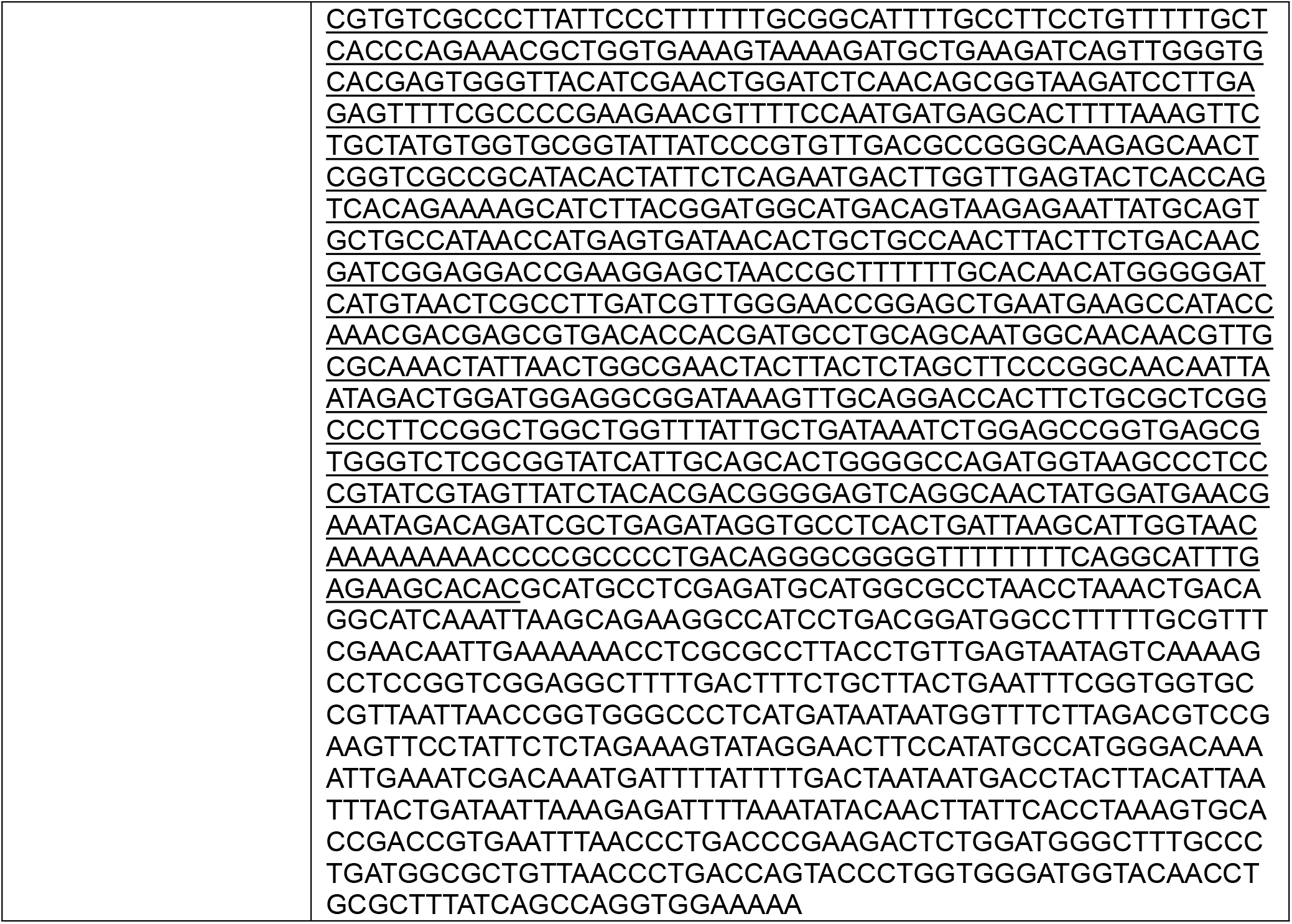
Sequence verification of genomic modifications to EcN.

**Figure S1:**
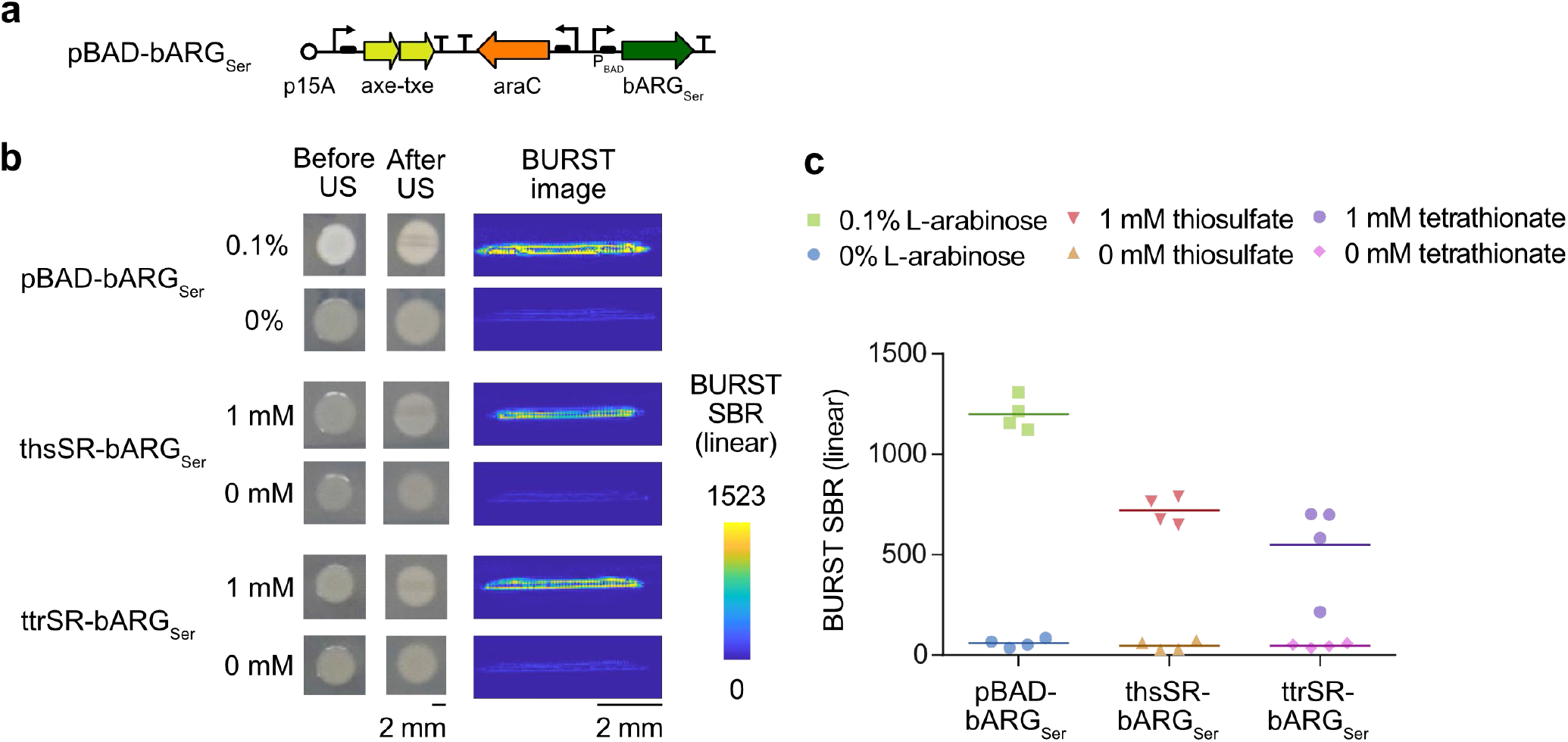
Arabinose-inducible versus initial thiosulfate and tetrathionate sensor constructs. (**a**) Plasmid diagram of the arabinose-inducible bARG_Ser_ construct pBAD-bARG_Ser_. (**b**) Images of patches of EcN strains containing the arabinose-inducible construct pBAD-bARG_Ser_, the initial thiosulfate sensor construct thsSR-bARG_Ser_, or the initial tetrathionate sensor construct ttrSR-bARG_Ser_ (see **Fig. 2a** for plasmid maps) on M9 plates without inducer or with 0.1% L-arabinose, 1 mM thiosulfate, or 1 mM tetrathionate, respectively. Photographs show the opacity of representative patches before ultrasound (US) imaging (left) or after being covered with agar and imaged using BURST ultrasound which reduces opacity along the imaging plane due to GV collapse (middle). Representative BURST images (right) show the ARG-specific ultrasound signal in cross-sections of the EcN patches. (**c**) Quantification of BURST images in terms of signal-to-background ratio (SBR).

**Figure S2:**
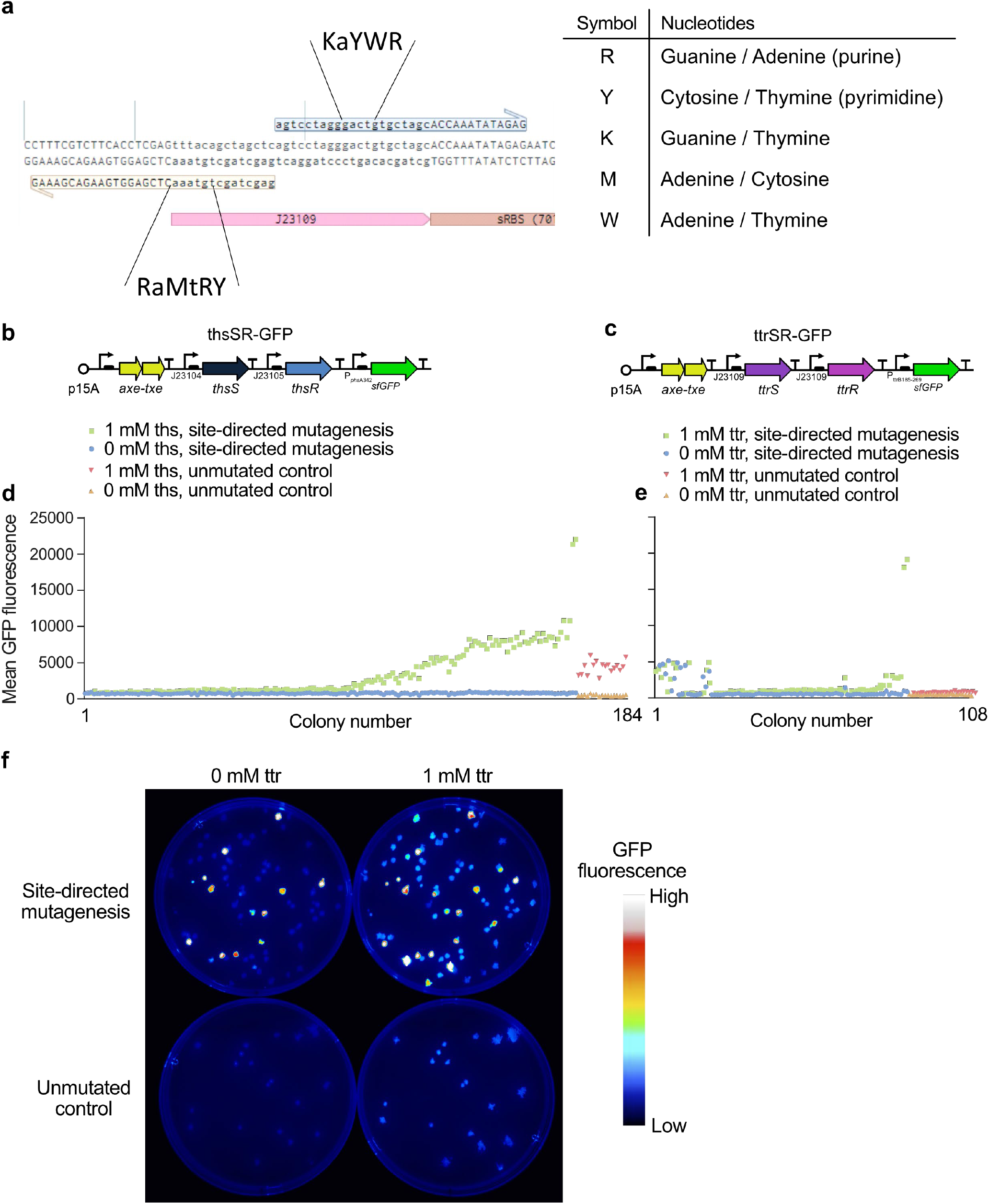
Site-directed mutagenesis and screening of GFP versions of thiosulfate and tetrathionate sensors. (**a**) Diagram of semi-random primers used for PCR for site-directed mutagenesis of the response regulator promoter, here J23109. The semi-random bases were chosen to broadly sample the Anderson promoter library^99^. (**b-c**) Plasmid diagrams of thiosulfate (thsSR-GFP) and tetrathionate (ttrSR-GFP) sensors with GFP as the output. (**d-e**) Mean GFP fluorescence of NEB Stable *E. coli* colonies from replica plating transformants on plates with 0 or 1 mM thiosulfate (ths) or tetrathionate (ttr) at 30°C after performing site-directed mutagenesis on the response regular promoter (site-directed mutagenesis) or leaving the plasmid unmutated (unmutated control). (d-e) Share the same y-axis values. (**f**) Representative GFP fluorescence images of replica plates of mutant library (top) or unmutated control (bottom) of ttrSR-GFP with 0 or 1 mM ttr.

**Figure S3:**
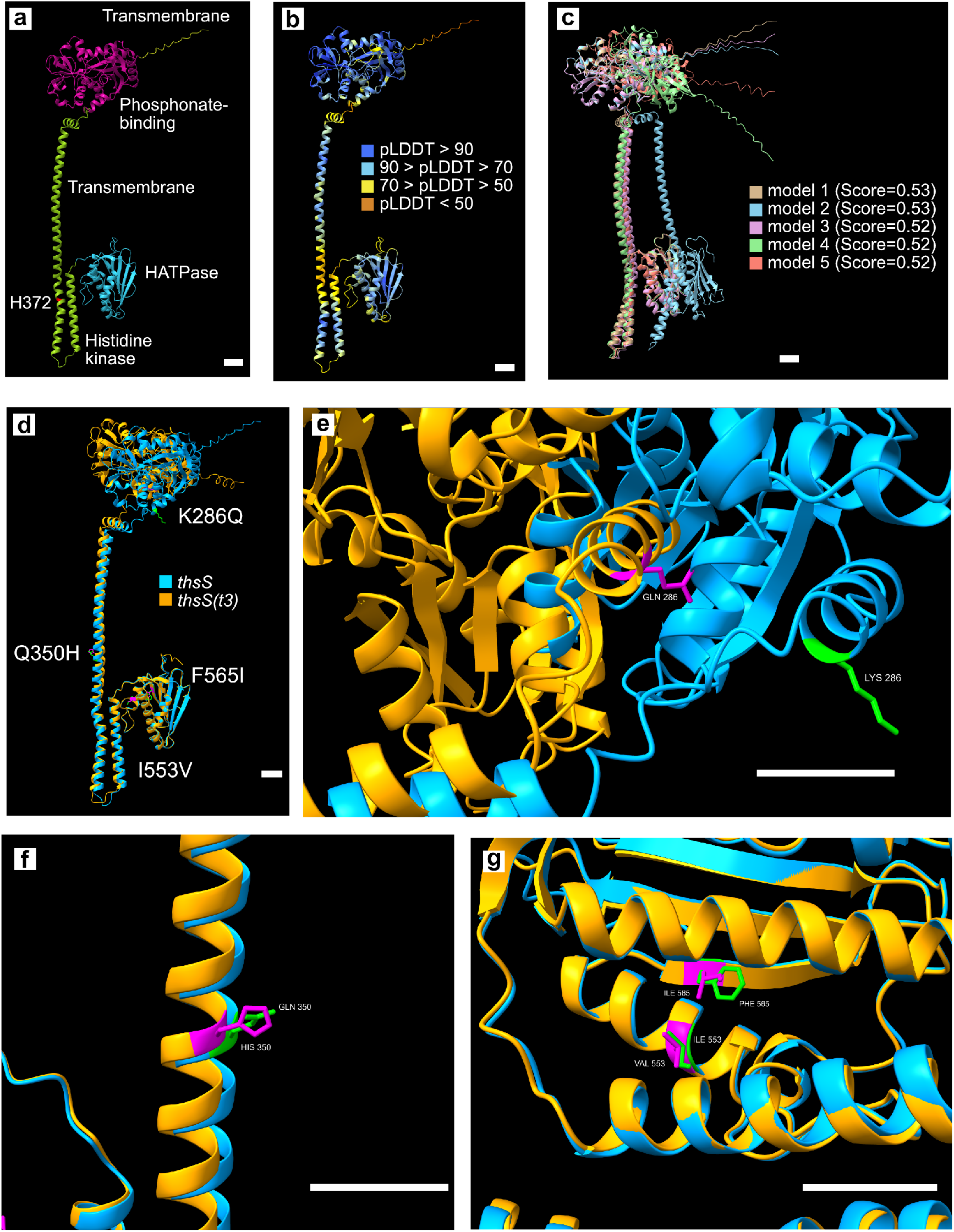
Structural predictions of the thiosulfate membrane sensor kinase protein *thsS* from *Shewanella halifaxensis* using AlphaFold 3^Ref. 97^. (**a**) Highest-ranked model of *thsS* colored by the predicted domains^24^. The histidine predicted to be involved in phospho-transfer is indicated in red (H372). (**b**) Highest-ranked model of *thsS* colored by the confidence score in terms of the pLDDT (predicted local distance difference test). (**c**) Alignment of the top-five highest ranked models of *thsS* and their corresponding overall ranking score. (**d**) Alignment of the highest-ranked models for *thsS* and the *thsS(t3)* variant which was identified in the screen depicted in **Fig. 2d**. The four amino acids that are different between the *thsS* and *thsS(t3)* are indicated in green and magenta, respectively. (**e-g**) Close-up views of the K286Q (e), Q350H (f), and I553V & F565I (g) point mutations from the structures depicted in (d). All scale bars are 10 Å.

**Figure S4:**
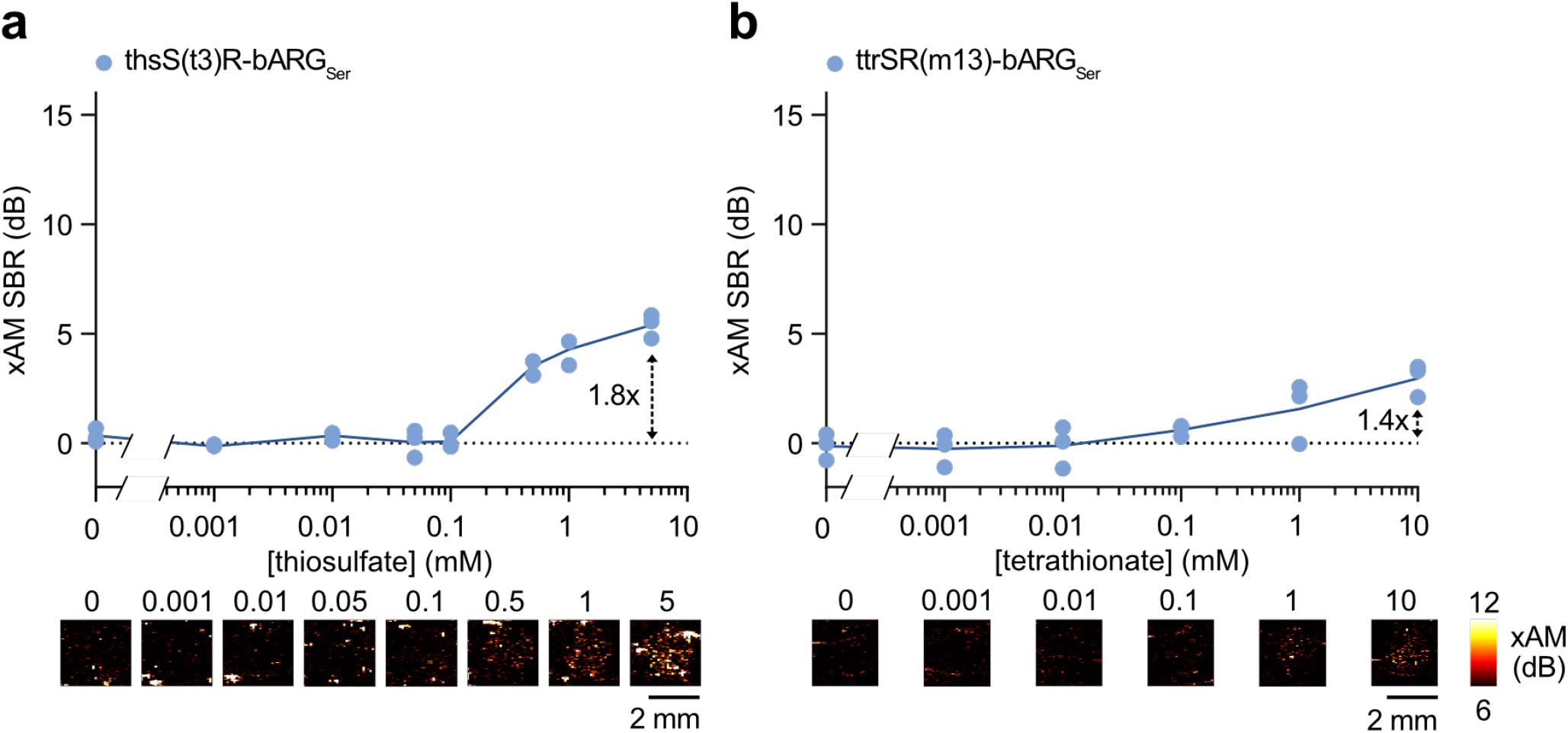
xAM ultrasound imaging of optimized thiosulfate and tetrathionate sensors. (**a-b**) Representative xAM ultrasound images (bottom) and quantification of the signal-to-background ratio (SBR) (top) of the best variants for thsSR-bARG_Ser_ (a) and ttrSR-bARG_Ser_ (b) at varying thiosulfate/tetrathionate concentrations in EcN at 37°C in liquid culture. Cells were cast in agarose phantoms at 10^9^ cells/mL for ultrasound imaging. Solid lines represent the mean of 3 biological replicates which are each averaged over 2 technical replicates.

**Figure S5:**
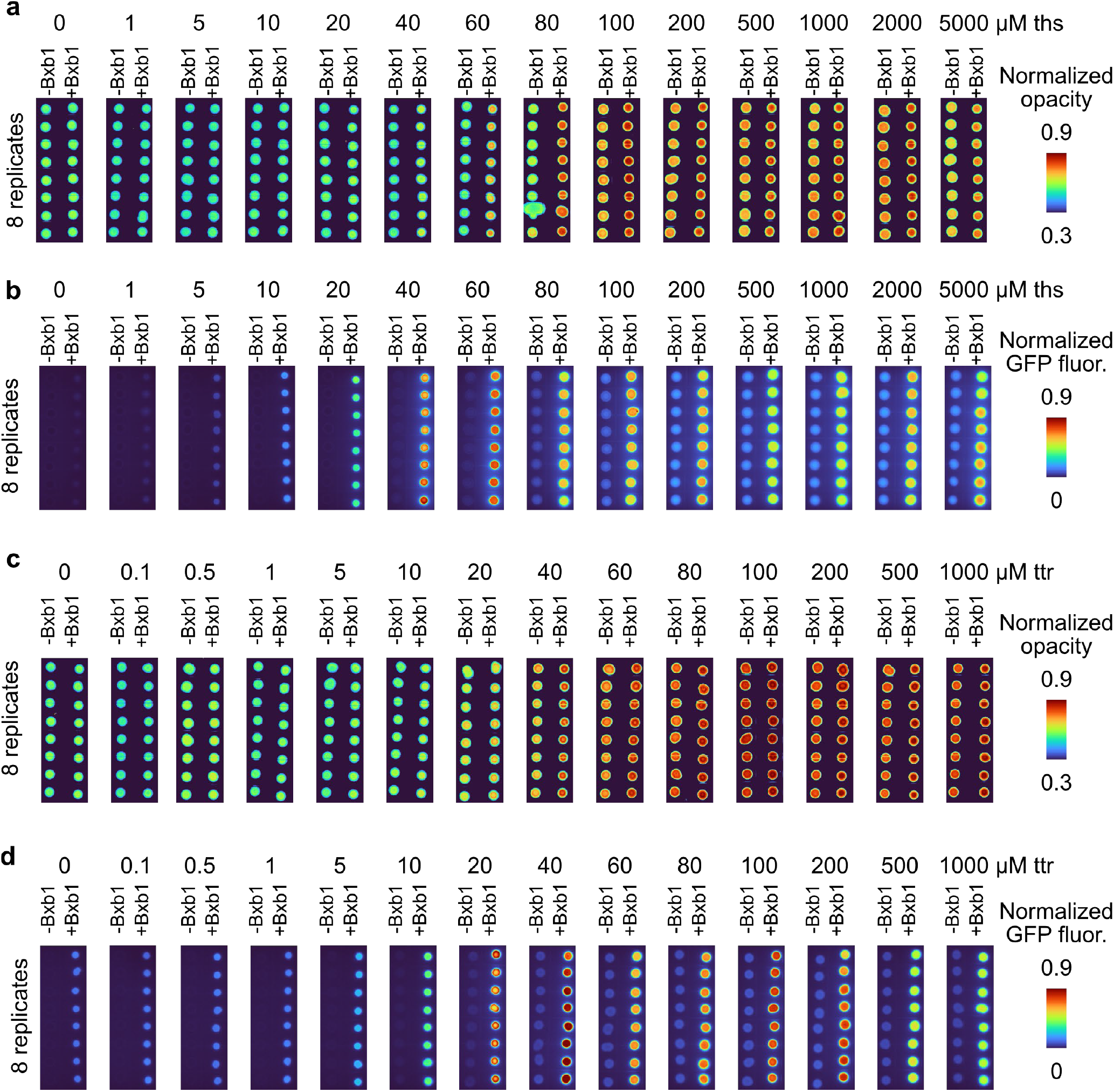
Images of patches of EcN sensor strains with and without the Bxb1 switch on plates with varying thiosulfate and tetrathionate concentrations. (**a**) Normalized transmitted white light images showing the opacity of thsS(t3)R-bARG_Ser_ (-Bxb1) and thsS(t3)R-Bxb1_P7-bARG_Ser_ (+Bxb1) EcN patches grown on plates at varying thiosulfate concentrations. (**b**) Normalized green fluorescence images of thsS(t3)R-GFP_mCherry (-Bxb1) and thsS(t3)R-Bxb1_P7-GFP_mCherry (+Bxb1) EcN patches grown on plates at varying thiosulfate concentrations. (**c**) Normalized transmitted white light images showing the opacity of ttrSR(m13)-bARG_Ser_ (-Bxb1) and ttrSR(m13)-Bxb1_P7-bARG_Ser_ (+Bxb1) EcN patches grown on plates at varying tetrathionate concentrations. (**d**) Normalized green fluorescence images of ttrSR(m13)-GFP_mCherry (-Bxb1) and ttrSR(m13)-Bxb1_P7-GFP_mCherry (+Bxb1) EcN patches grown on plates at varying tetrathionate concentrations. All patches were grown on M9 plates at 37°C. Patches were suspended in PBS for the ultrasound imaging and flow cytometry depicted in **Fig. 3 and S6**.

**Figure S6:**
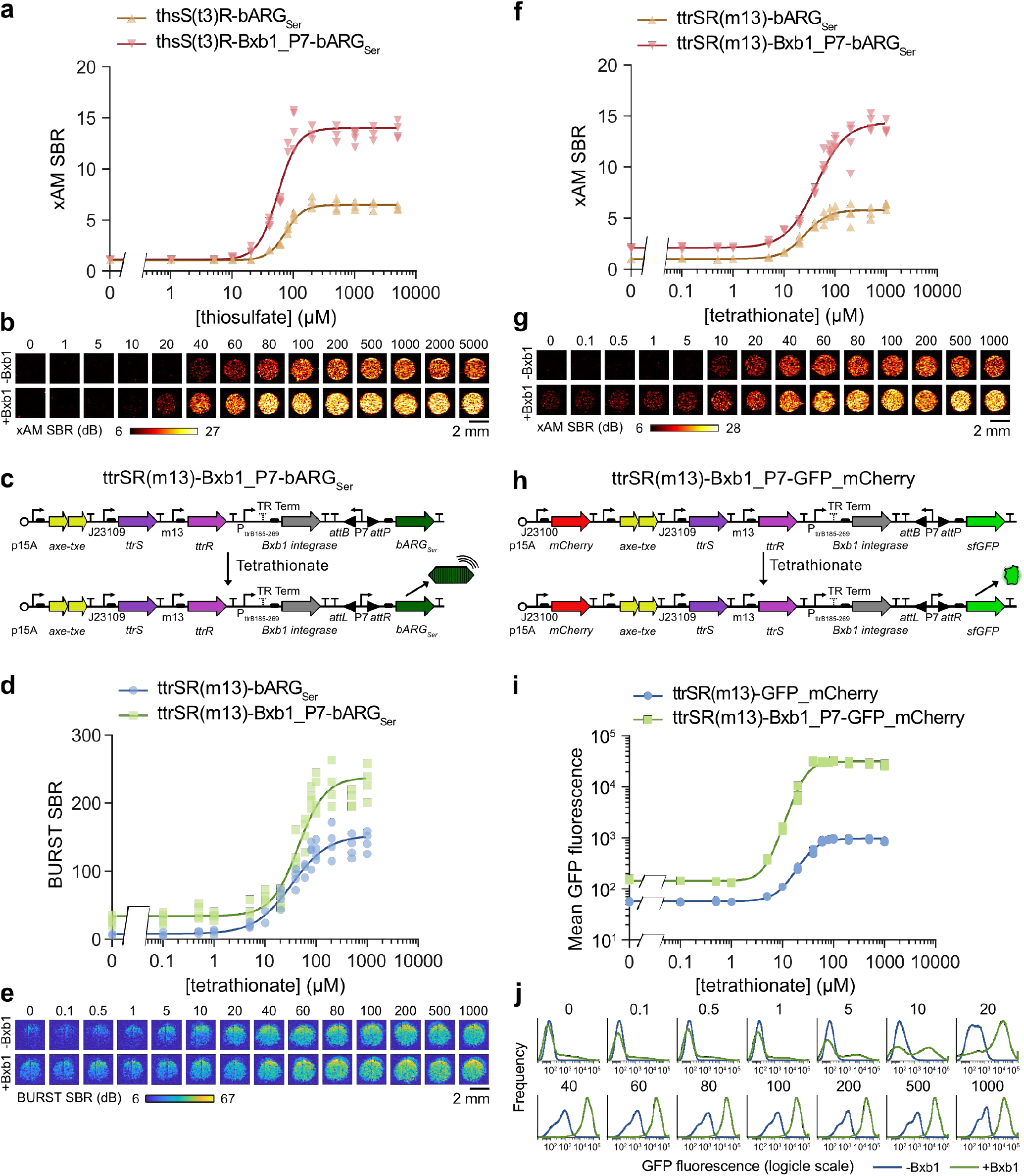
Additional in vitro characterization data for integrase-based switch sensors. (**a-b**) xAM signal-to-background ratio (SBR) (a) and representative images (b) of the optimized thiosulfate sensor with and without the Bxb1 integrase-based switch at varying thiosulfate concentrations. See **Fig. 3b-c** for the corresponding BURST data. (**c**) Plasmid diagram of the optimized tetrathionate sensor ttrSR(m13)-bARG_Ser_ with an integrase-based switch to create ttrSR(m13)-Bxb1_P7-bARG_Ser_. (**d-e**) BURST signal-to-background ratio (SBR) (d) and representative images (e) of the optimized tetrathionate sensor with and without the Bxb1 integrase-based switch at varying tetrathionate concentrations. (**f-g**) xAM signal-to-background ratio (SBR) (f) and representative images (g) of the optimized tetrathionate sensor with and without the Bxb1 integrase-based switch at varying tetrathionate concentrations. (**h**) Plasmid diagram of the optimized tetrathionate sensor ttrSR(m13)-GFP_mCherry with an integrase-based switch to create ttrSR(m13)-Bxb1_P7-GFP_mCherry. (**i-j**) Mean GFP fluorescence measured via flow cytometry (i) and representative histograms (j) of the optimized tetrathionate sensor with and without the Bxb1 integrase-based switch at varying tetrathionate concentrations. In (a), (d), (f), and (i), points represent biological replicates (N=4) and curves represent fits to the Hill equation (see **Table S2** for fitted parameters). All strains were grown on plates with varying concentrations of thiosulfate and tetrathionate at 37°C (see **Fig. S5** for images of the plates) and suspended in PBS for ultrasound imaging and flow cytometry; for ultrasound imaging, cells were cast in agarose phantoms at a concentration of 5 × 10^8^ cells/mL.

**Figure S7:**
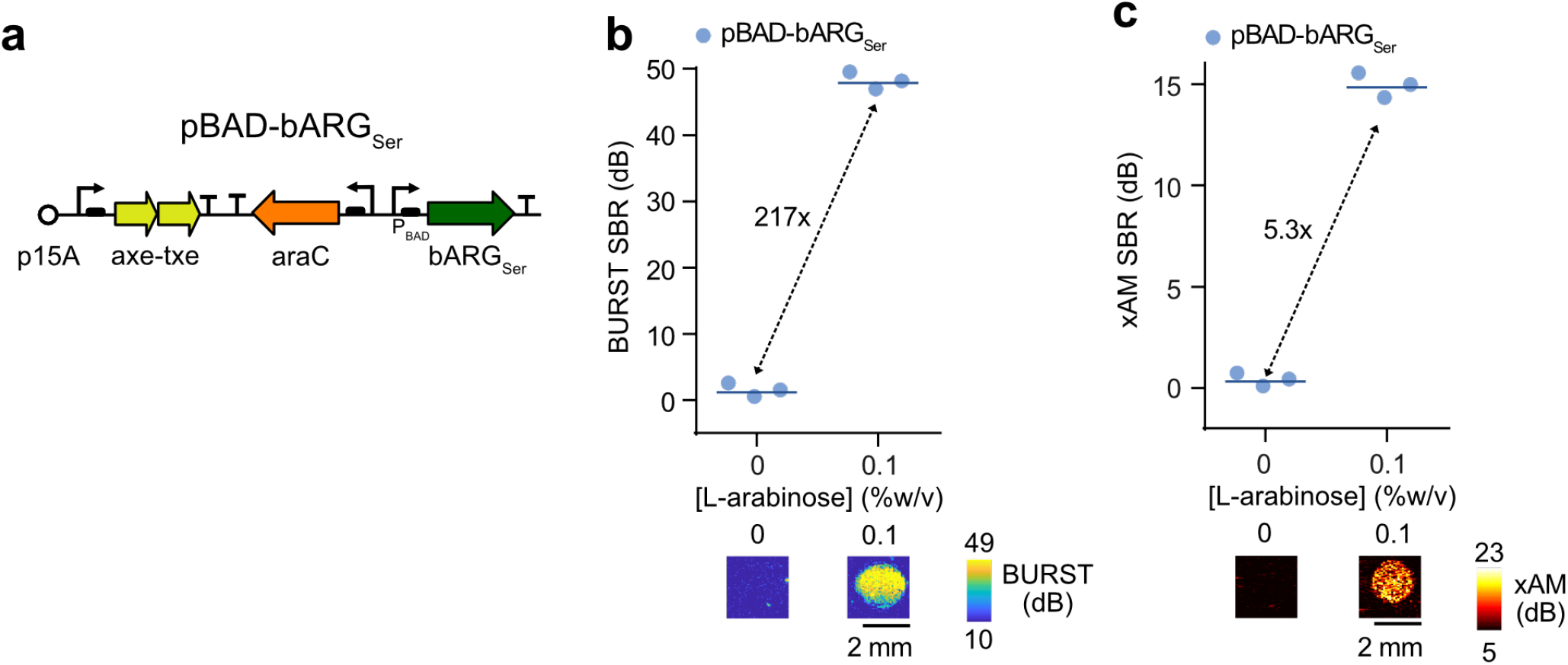
In vitro characterizations of arabinose-inducible bARG_Ser_ in EcN. (**a**) Plasmid diagram of the arabinose-inducible bARG_Ser_ construct pBAD-bARG_Ser_. (**b-c**) Representative BURST (b) and xAM (c) ultrasound images (bottom) and quantification of the signal-to-background ratio (SBR) (top) of pBAD-bARG_Ser_ at 0 and 0.1% L-arabinose in *E. coli* Nissle (EcN) at 37°C in liquid culture. Cells were cast in agarose phantoms at 10^9^ cells/mL for ultrasound imaging. Lines represent the mean of 3 biological replicates which are each averaged over 2 technical replicates.

**Figure S8:**
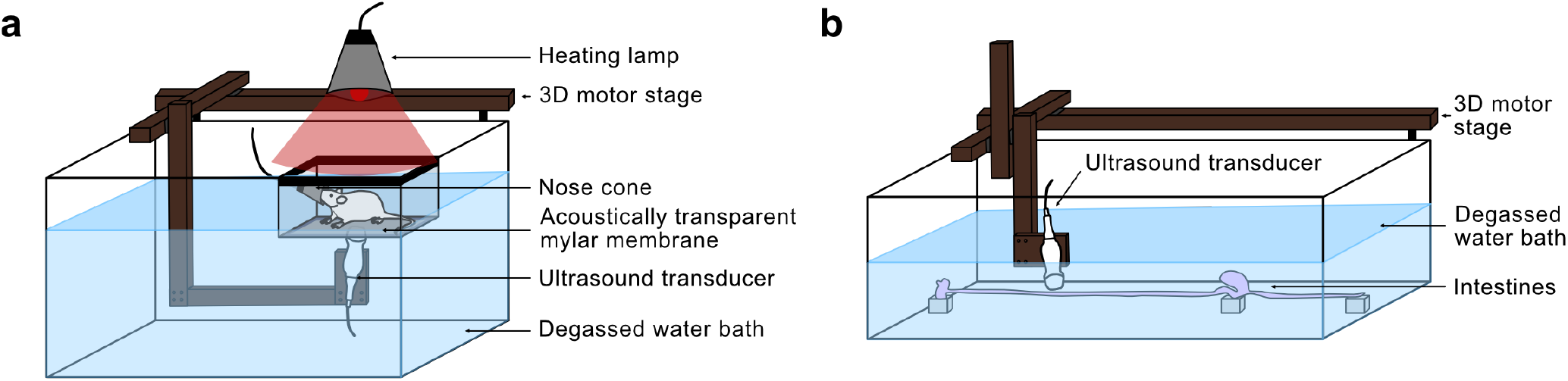
Custom-built in vivo and ex vivo ultrasound scanning systems. (**a-b**) Diagrams of the ultrasound imaging setups for fast and easy scanning of the entire abdominal area of live mice (a) and of the entire mouse GI tract ex vivo (b).

**Figure S9:**
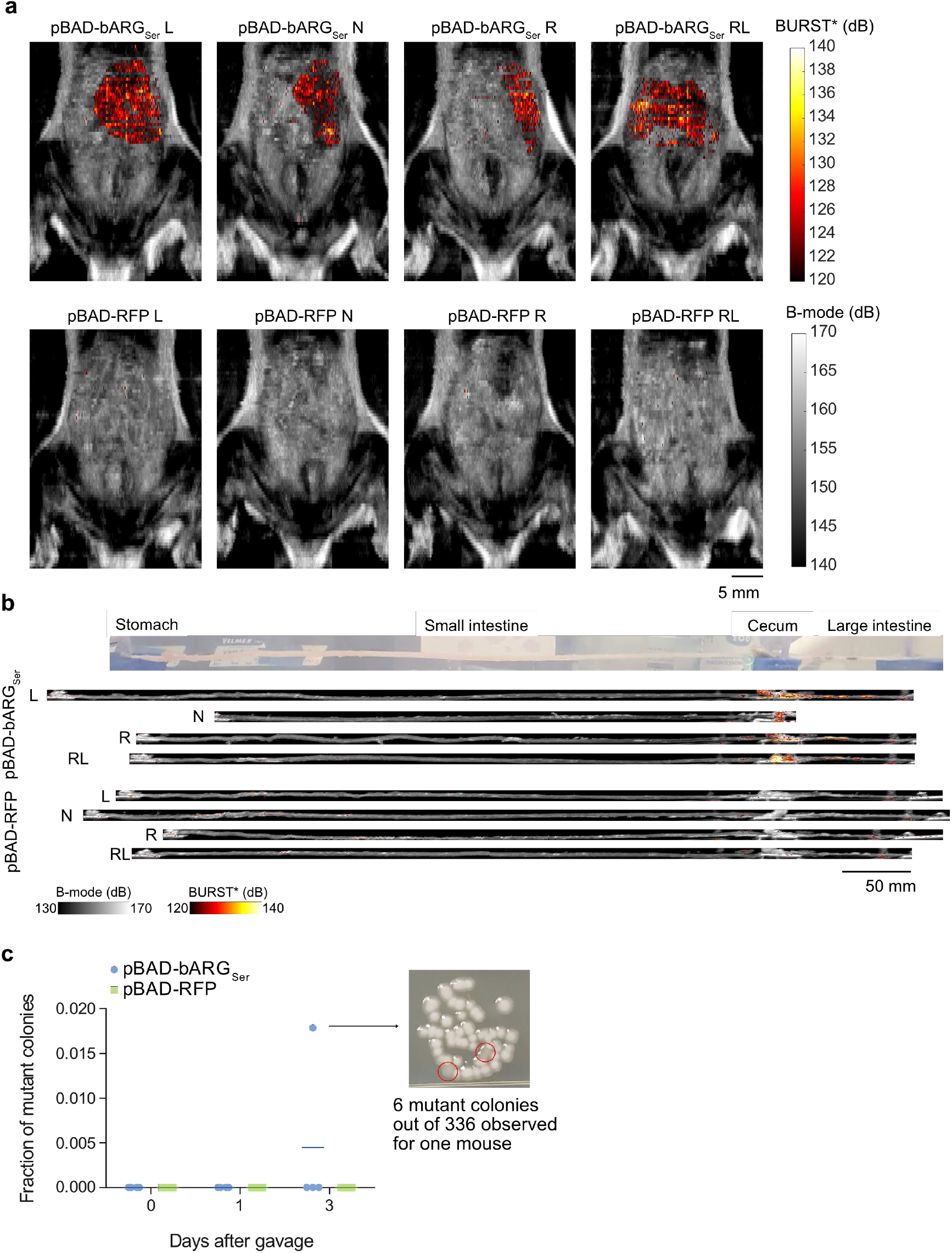
All replicates and additional data for imaging arabinose-inducible bARG_Ser_ expression in EcN colonizing the GI tract. (**a**) Ultrasound images overlaying the integrated BURST* signal over the depth onto the integrated B-mode signal over the depth for all mice one day after administration of the L-arabinose-sensing EcN using the setup depicted in **Fig. S8a**. (**b**) Ex vivo ultrasound images of intestines from all mice 3 days after administration of the L-arabinose-sensing EcN using the setup depicted in **Fig. S8b**. The integrated BURST* signal over the width was overlaid onto the integrated B-mode signal over the width. (**c**) Fraction of non-opaque or non-RFP-fluorescent mutant colonies detected by plating the gavage mixtures (day 0) or the feces on days 1 and 3 after gavage of the L-arabinose-sensing EcN onto plates with L-arabinose. Six non-opaque colonies (out of 336 total colonies) were observed in one mouse colonized by pBAD-bARG_Ser_ EcN on day 3. Three of these colonies (red circles) are depicted in the image on the right.

**Figure S10:**
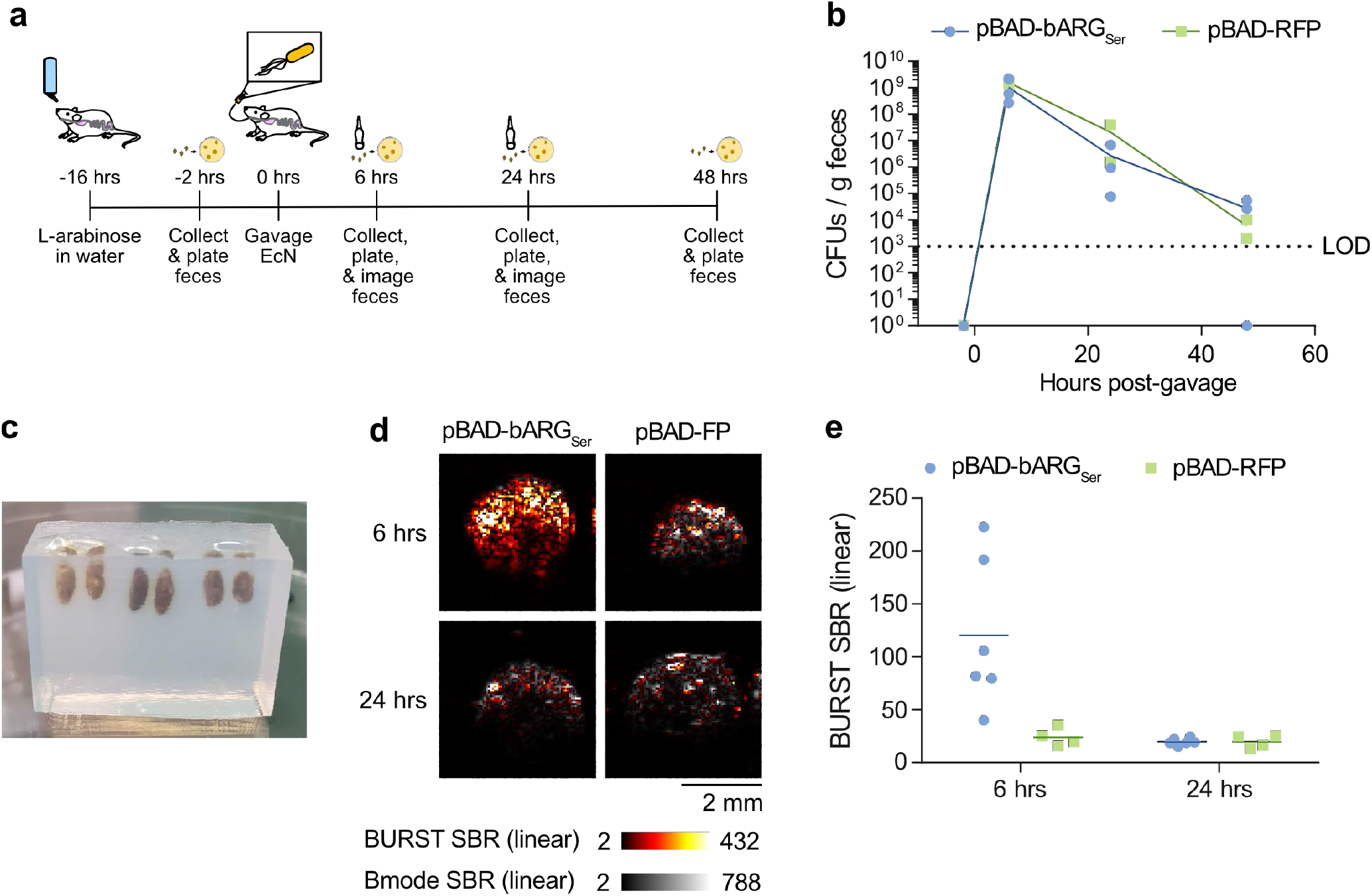
Ultrasound imaging of bARG_Ser_-expressing EcN in feces without antibiotic treatment. (**a**) Experimental design for testing L-arabinose-inducible bARG_Ser_ expression in EcN in vivo without antibiotics. Mice were given water containing L-arabinose for 16 hours, and EcN containing pBAD-bARG_Ser_ or the control plasmid pBAD-RFP were orally administered. Feces were collected at various time points and imaged with ultrasound and/or plated on selective media to measure colonization. (**b**) Colony forming units (CFUs) per gram of feces collected 2 hours before, and 6, 24, and 48 hours after oral gavage of the EcN strains. Limit of detection (LOD) was 1.7 × 10^3^ CFU/g feces. N = 3 mice per strain. (**c-e**) Representative phantom containing feces for ultrasound imaging (c), representative ultrasound images of feces overlaying the thresholded BURST image (hot scale) over the B-mode image (grayscale) (d), and quantification of the BURST signal-to-background ratio (SBR) of feces (e). Two fecal pellets were imaged per mouse, giving N = 6 per strain. In (b) and (e), points represent biological replicates and lines represent the mean.

**Figure S11:**
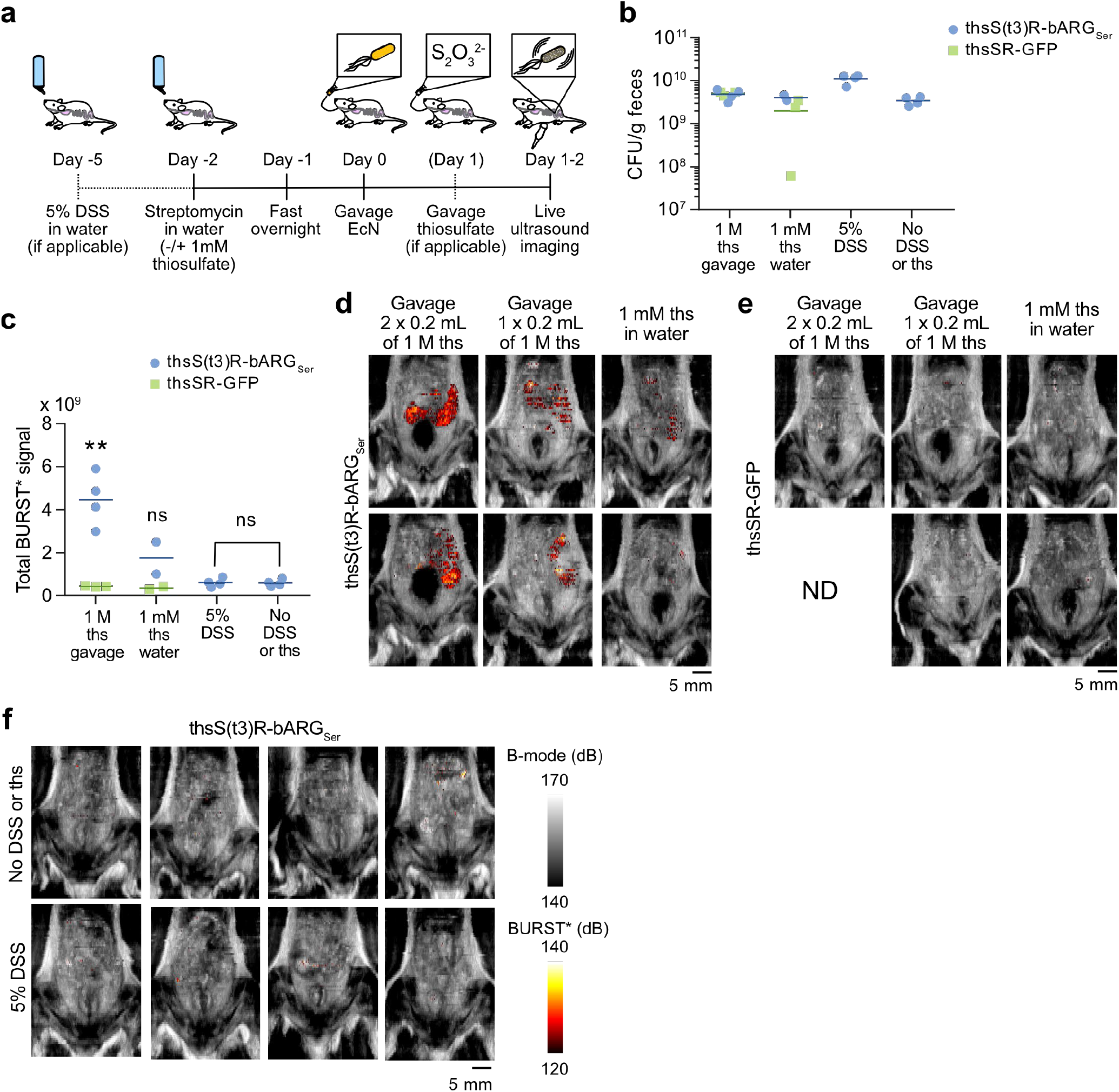
All replicates and additional data for testing thsS(t3)R-bARG_Ser_ and thsSR-GFP sensor EcN strains in vivo. (**a**) Diagram summarizing the experimental design involving DSS treatment, thiosulfate in the drinking water, or thiosulfate administered via oral gavage. All mice received streptomycin in the drinking water for 2 days before oral gavage of sensor EcN strains, and all mice were imaged using the setup depicted in **Fig. S8a** one to two days after EcN gavage. (**b**) Colony forming units (CFU) per gram of feces one day after oral gavage of the sensor EcN strains. (**c**) Total BURST* ultrasound signal imaged in mice one day (1 mM ths water, 5% DSS, and no DSS or ths) or two days (1 M ths gavage) after EcN gavage. (**d-e**) Overlay of the integrated BURST* images onto the integrated B-mode images over the depth for all mice colonized by thsS(t3)R-bARG_Ser_ (d) or thsSR-GFP (e) EcN and treated with thiosulfate via oral gavage or in the drinking water. (**f**) Overlay of the integrated BURST* images onto the integrated B-mode images over the depth for all mice colonized by thsS(t3)R-bARG_Ser_ EcN and treated with DSS or left untreated (no DSS or thiosulfate). Asterisks represent statistical significance by unpaired Student’s t-tests (** = p<0.01, ns = no significance).

**Figure S12:**
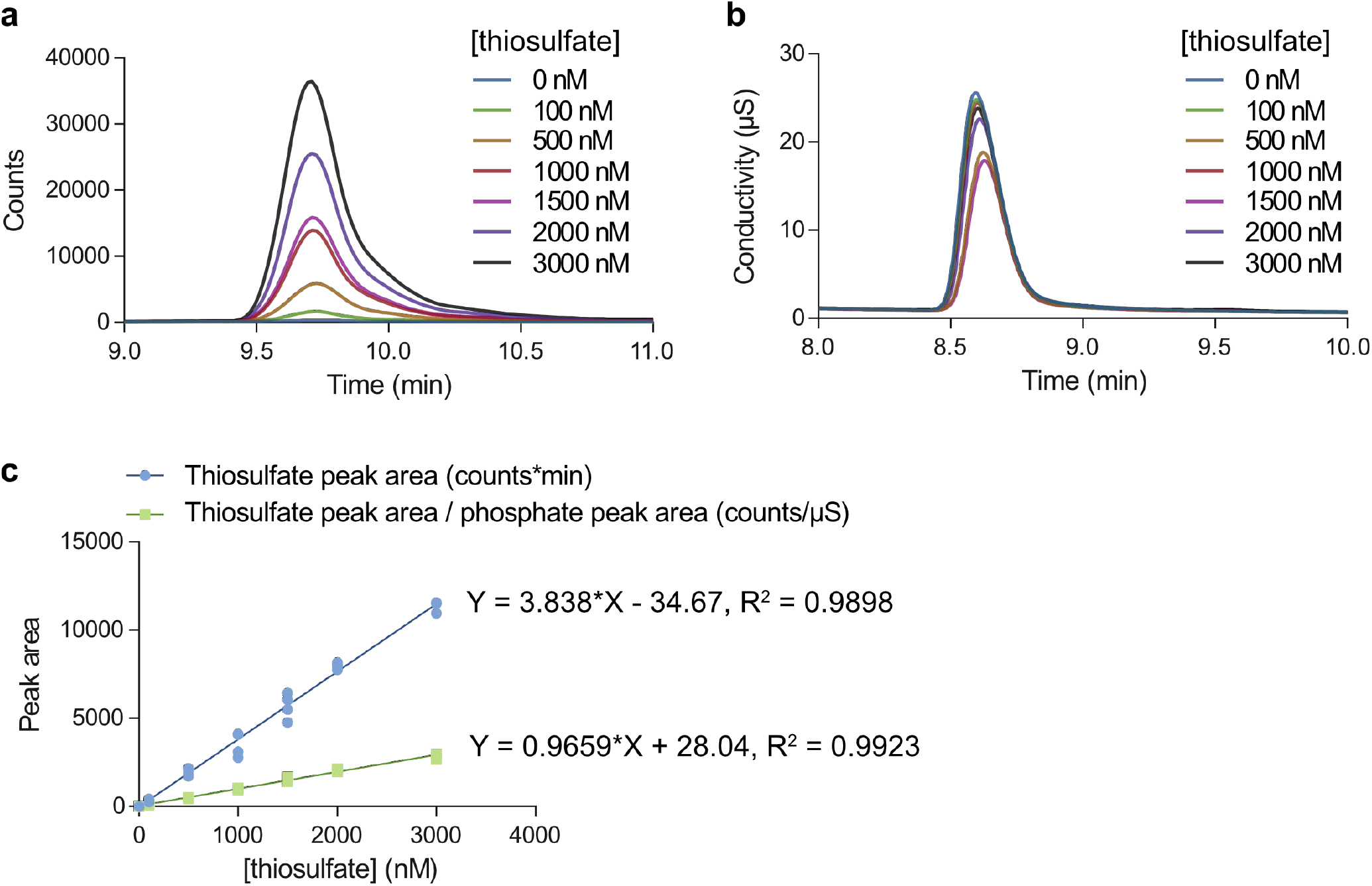
Ion chromatography-mass spectrometry (IC-MS) chromatograms and standard curves for quantifying thiosulfate. (**a**) Representative extracted ion chromatograms showing the thiosulfate peaks (m/z = 112.5-113.5 filter) in thiosulfate samples at concentrations ranging from 0 to 3000 nM in a PBS background. (**b**) Representative conductivity chromatograms showing the phosphate peaks in thiosulfate samples at concentrations ranging from 0 to 3000 nM in a PBS background. The concentration of PBS was the same in all samples, so the phosphate served as an internal standard to correct for variations in the injection volume. (**c**) Standard curve for the raw thiosulfate peak area and for the thiosulfate peak area normalized by the corresponding phosphate peak area. The normalized peak areas were used for quantification of thiosulfate in fecal and intestinal samples because normalization resulted in less variation between technical replicates. Points represent technical replicates (N=4) and lines represent a linear regression.

**Figure S13:**
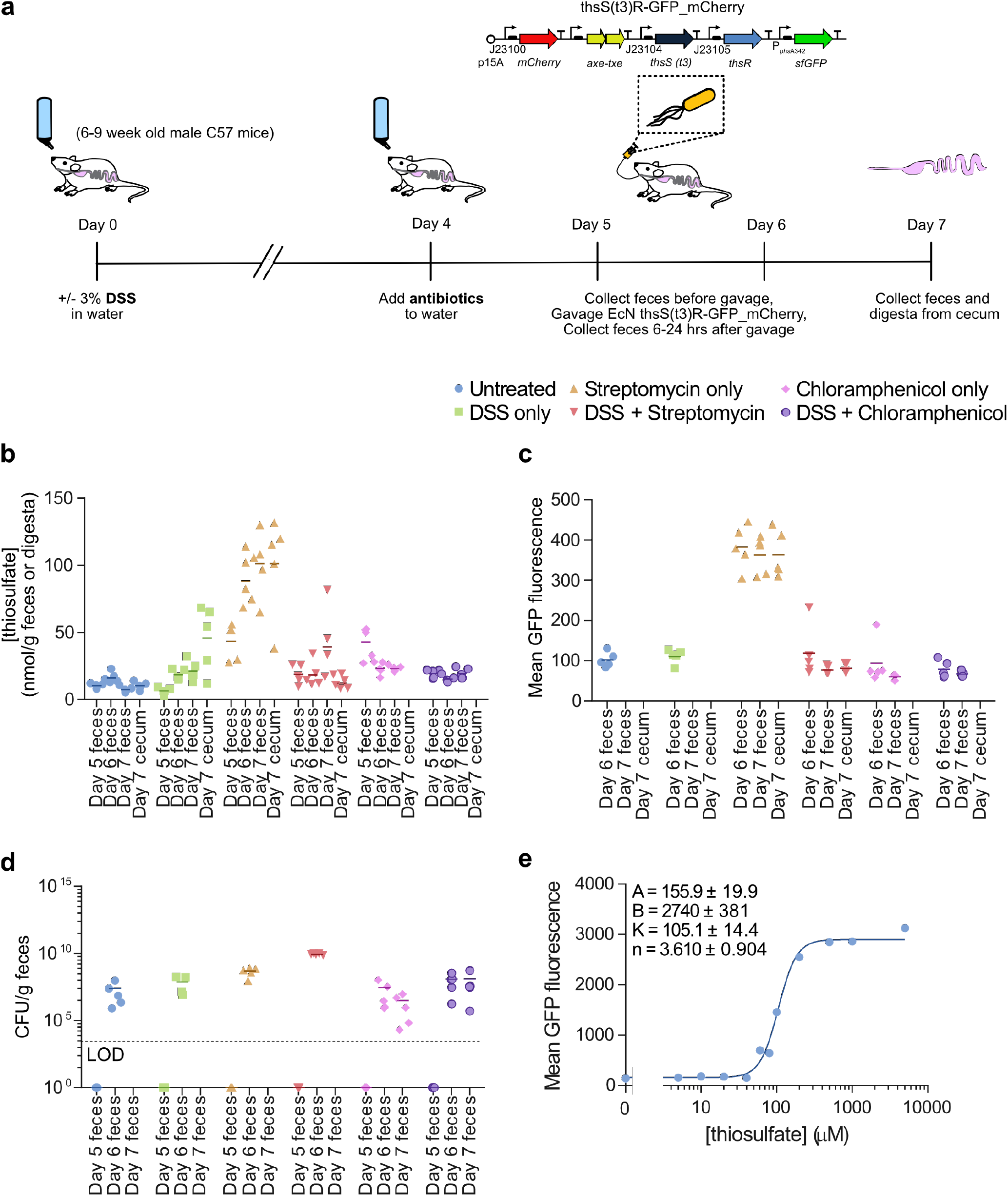
Measuring thiosulfate levels and thsS(t3)R-GFP_mCherry sensor activation in response to DSS and antibiotics in vivo. (**a**) Experimental design for testing the effect of DSS and antibiotics on intestinal thiosulfate levels and thiosulfate sensor activation. Mice were given water with 3% DSS or without DSS on day 0, and on day 4 antibiotics were added to the water. The thiosulfate-sensing EcN strain with plasmid thsS(t3)R-GFP_mCherry was administered via oral gavage on day 5 for antibiotic-treated mice and on day 6 for mice not given antibiotics. Feces were collected on days 5-7, cecal contents were collected on day 7, and they were analyzed via IC-MS, flow cytometry, and plating. (**b**) Concentrations of thiosulfate measured via IC-MS in the feces or digesta of the cecum. (**c**) Mean GFP fluorescence of positive mCherry events of the sensor strain measured via flow cytometry. There is no flow cytometry data for mice not treated with antibiotic on day 7 because the strain did not colonize without antibiotics. (**d**) Colony forming units (CFU) per gram of feces measured by plating on selective media to assess colonization of the sensor bacteria. (**e**) In vitro characterization of the thsS(t3)R-GFP_mCherry EcN sensor strain in terms of mean GFP fluorescence measured via flow cytometry after inducing with varying thiosulfate concentrations in liquid culture at 37°C. Maximal sensor activation observed in vivo in streptomycin-treated mice was only 11.8% that of maximal sensor activation observed in vitro.

**Figure S14:**
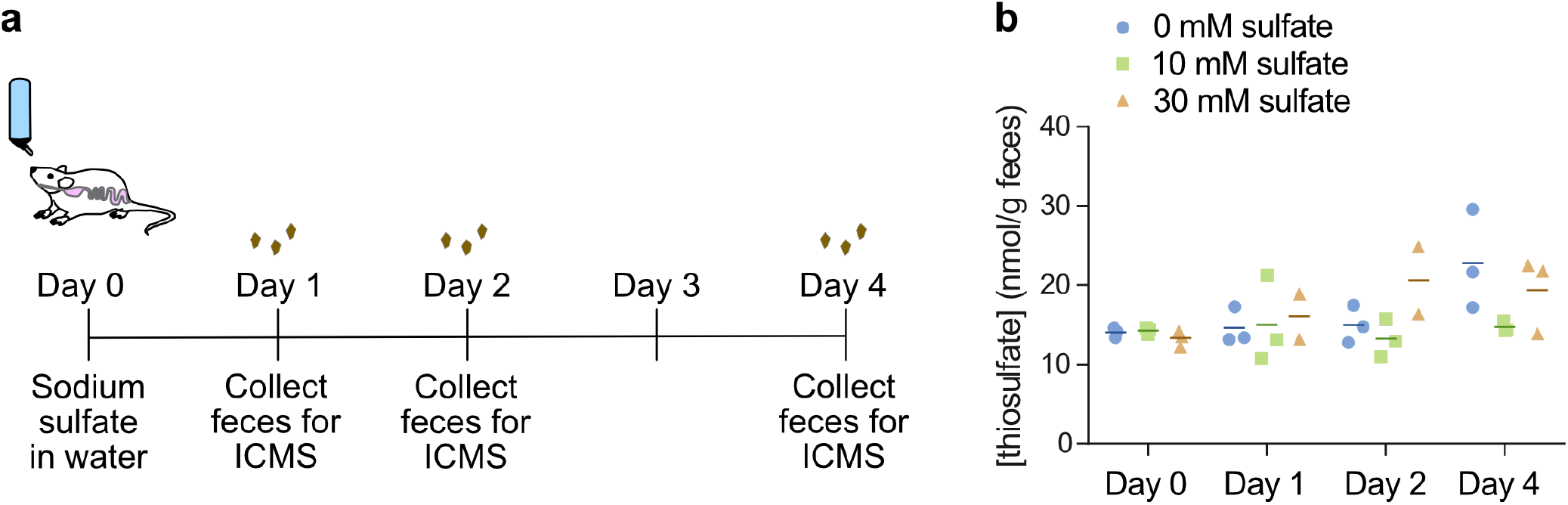
Effect of sodium sulfate in drinking water on fecal thiosulfate levels. (**a**) Experimental design, where mice were given drinking water containing 0, 10, or 30 mM sodium thiosulfate on day 0, and feces were collected and analyzed via IC-MS on days 0, 1, 2, and 4. 10 mM sodium thiosulfate corresponds to the concentration of sulfate in 5 g/L streptomycin sulfate. (**b**) Concentration of fecal thiosulfate measured by IC-MS. Supplementing the drinking water with sulfate did not significantly affect the fecal thiosulfate levels at any time point. Points represent biological replicates (N = 3 mice) and lines represent the mean.

**Figure S15:**
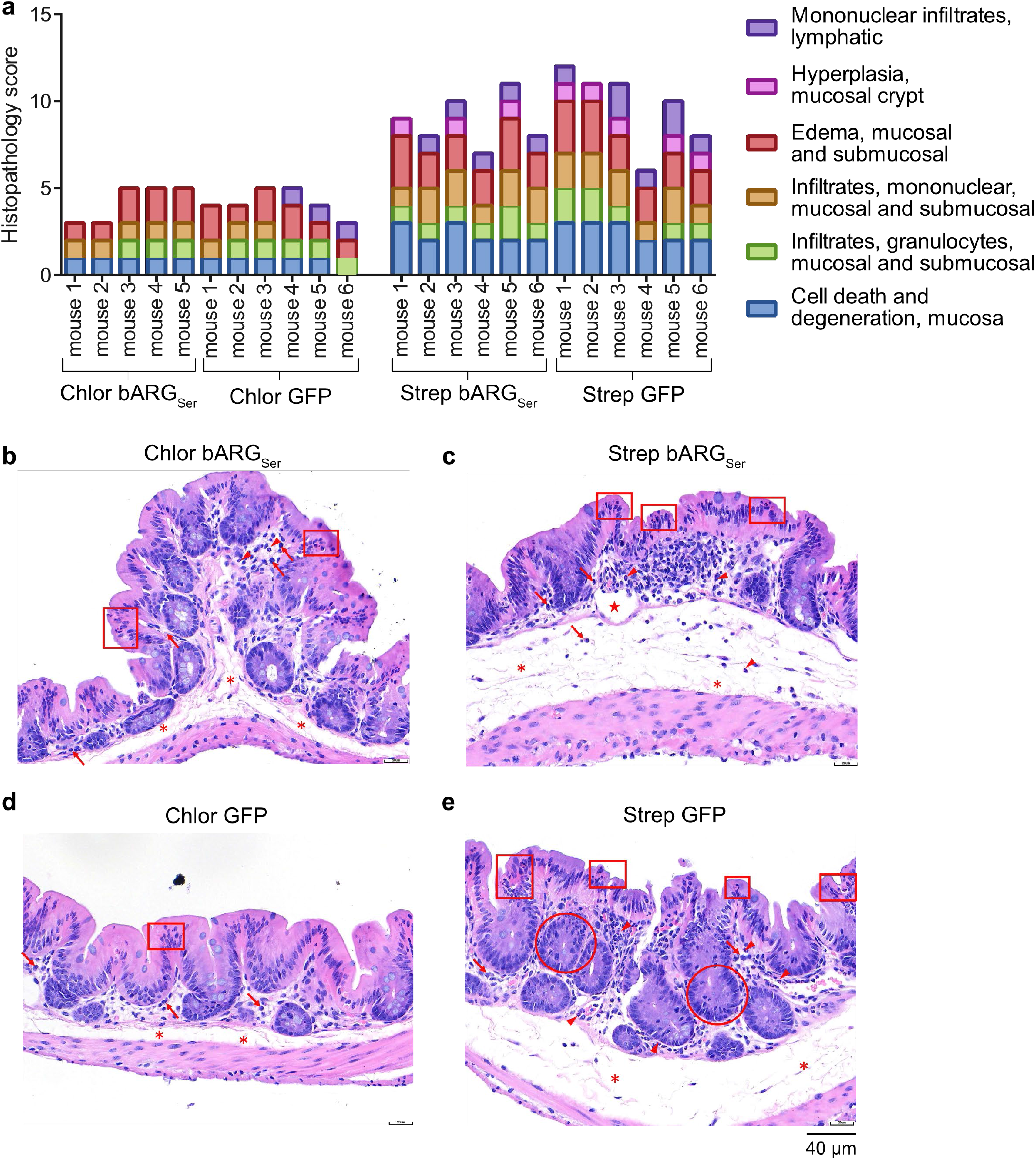
Full scoring data and additional histopathology images of cecal tissues from chloramphenicol- and streptomycin-treated mice. (**a**) Histopathology scoring of cecal tissues from chloramphenicol- and streptomycin-treated mice on day 5 of treatment broken down by category of abnormality. These data were aggregated for display in **Fig. 5d**. (**b-e**) Representative images of H&E-stained sections of cecal tissue on day 5 of antibiotic treatment for mice treated with chloramphenicol and colonized with thsS(t3)R-Bxb1_P7-bARG_Ser_ EcN (b), treated with streptomycin and colonized with thsS(t3)R-Bxb1_P7-bARG_Ser_ EcN (c), treated with chloramphenicol and colonized with thsS(t3)R-Bxb1_P7-GFP_mCherry EcN (d), and treated with streptomycin and colonized with thsS(t3)R-Bxb1_P7-GFP_mCherry EcN (e). Abnormalities are indicated in red: mucosal epithelial cell death and degeneration (box), mucosal crypt hyperplasia (circle), mucosal/submucosal edema (asterisk), mononuclear infiltrates (arrow), granulocytic infiltrates (arrowhead), dilated lymphatic (star).

**Figure S16:**
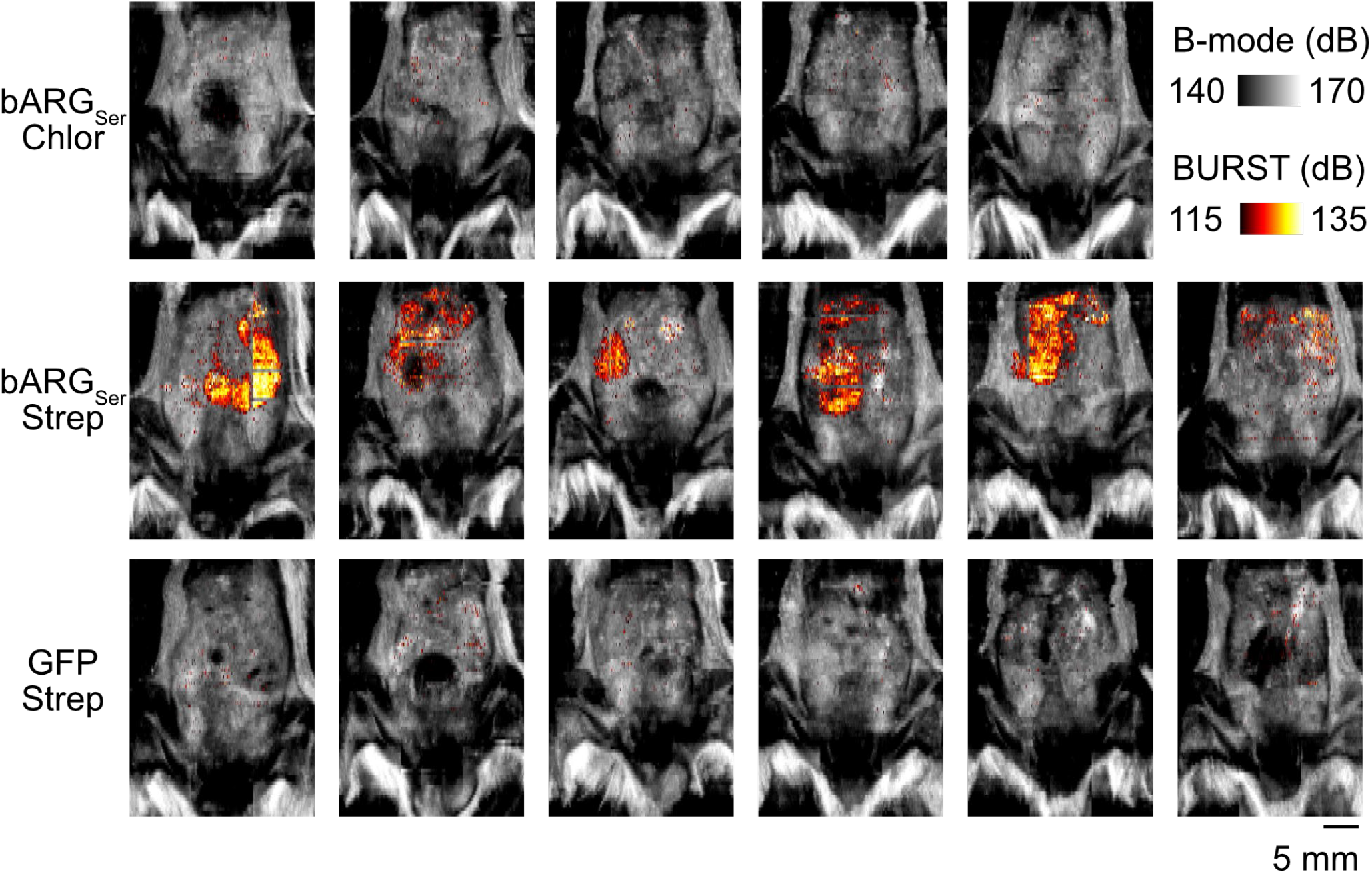
All ultrasound images of mice from the experiment depicted in Fig. 5. The integrated BURST* signal was overlaid onto the integrated B-mode signal for all mice treated with chloramphenicol and colonized with thsS(t3)R-Bxb1_P7-bARG_Ser_ EcN (bARG_Ser_ Chlor), for all mice treated with streptomycin and colonized with thsS(t3)R-Bxb1_P7-bARG_Ser_ EcN (bARG_Ser_ Strep), and for all mice treated with streptomycin and colonized with thsS(t3)R-Bxb1_P7-GFP_mCherry EcN (GFP Strep) on day 3.

**Figure S17:**
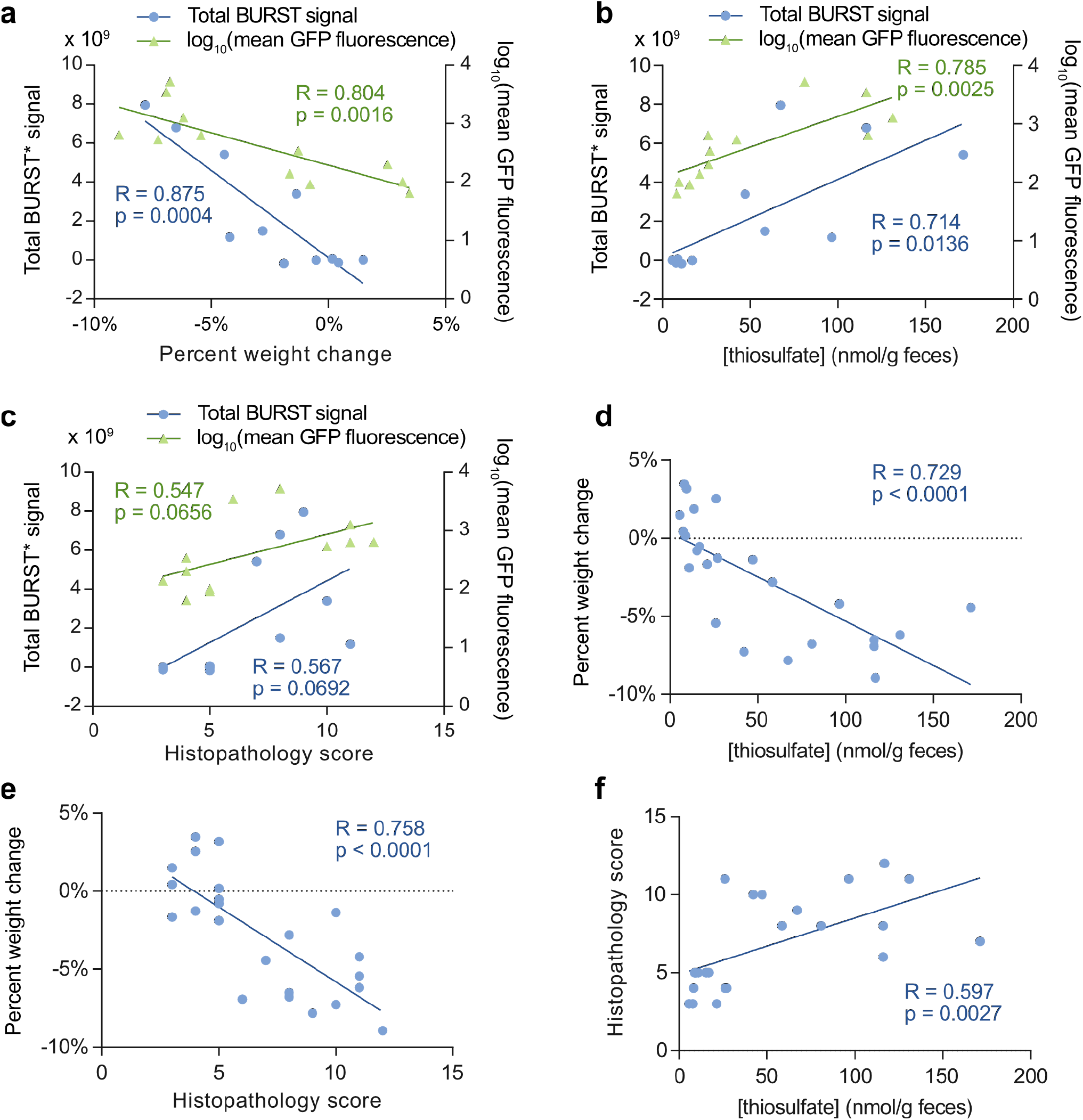
Correlations between thiosulfate sensor activation and disease severity in chloramphenicol- and streptomycin-treated mice. (**a-c**) Total BURST* signal imaged on day 3 or mean GFP fluorescence measured on day 4 versus the percent weight change on day 2 before mice were fasted with tail cups (a), versus the fecal thiosulfate levels measured via IC-MS on day 4 (b), and versus the histopathology score of cecal tissue on day 5 (c). (**d-e**) percent weight change on day 2 before mice were fasted with tail cups versus the fecal thiosulfate levels measured via IC-MS on day 4 (d), and versus the histopathology score of cecal tissue on day 5 (e). (**f**) Histopathology score of cecal tissue on day 5 versus the fecal thiosulfate levels measured via IC-MS on day 4. Lines represent linear regressions where R represents the goodness of fit and the p value indicates whether the slope is significantly non-zero.

**Figure S18:**
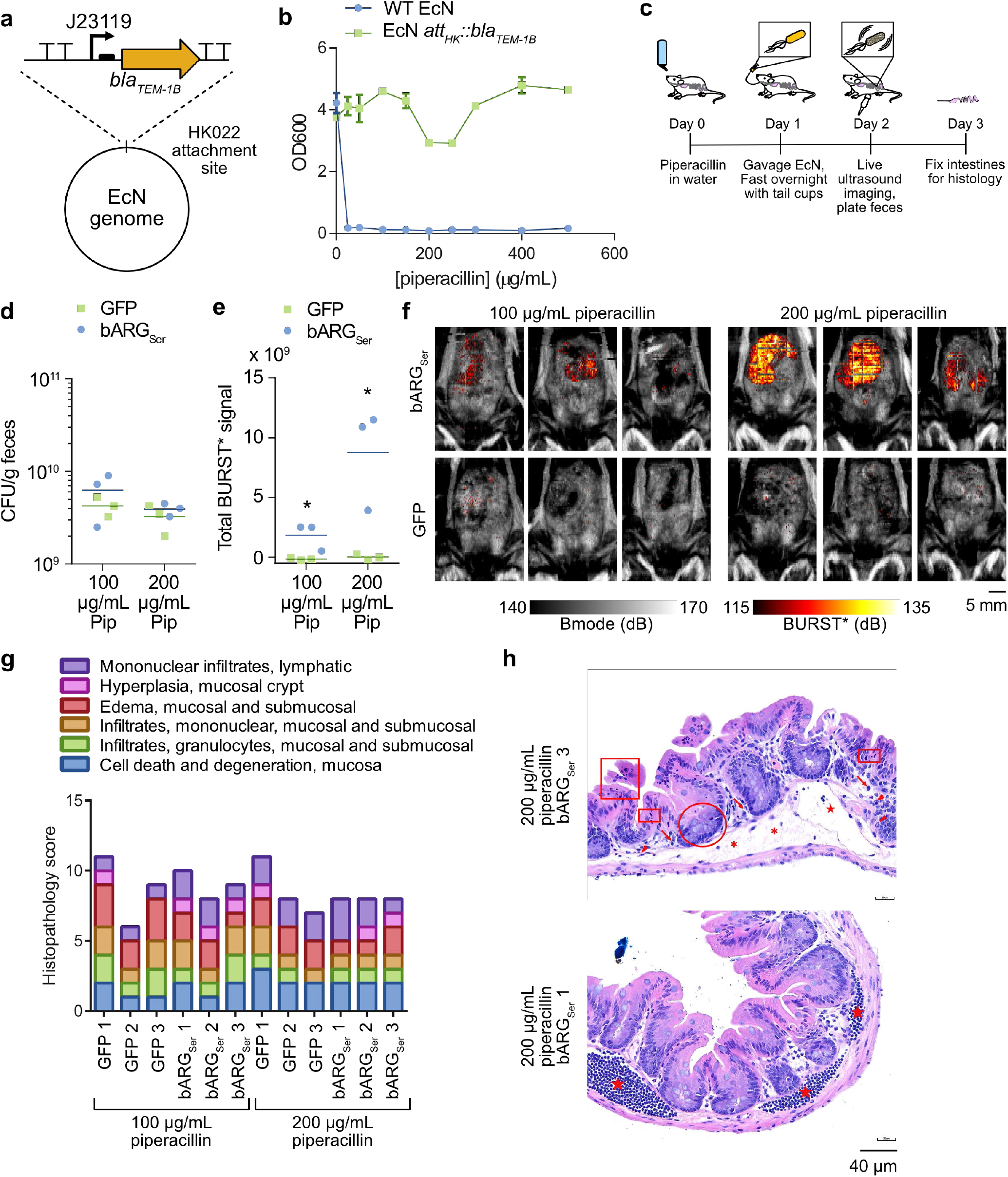
Ultrasound imaging of thiosulfate sensor activation in piperacillin-treated mice. (**a**) Diagram of genomic modification of EcN to confer piperacillin resistance. The beta lactamase gene *bla*_*TEM-1B*_ known to confer piperacillin resistance in *E. coli*^100,101^ was placed under control of the strong constitutive promoter J23119. This cassette was integrated into the genome at the phage HK022 attachment site in EcN using the clontegration system^102^. See **Table S3** for sequencing confirmation. (**b**) Optical density at 600 nm (OD600) after incubating wild-type (WT) EcN and the att_HK_:bla_TEM-1B_ EcN in media at varying piperacillin concentrations to confirm piperacillin resistance of the att_HK_:bla_TEM-1B_ strain. Points represent the mean of two biological replicates, error bars represent the standard deviation, and lines connect the points. (**c**) Experimental design for testing EcN strains containing plasmids for the optimized integrase-based switch thiosulfate sensors, thsS(t3)R-Bxb1_P7-bARG_Ser_ or thsS(t3)R-Bxb1_P7-GFP_mCherry, in piperacillin-treated mice. One day after piperacillin was administered via drinking water, the EcN strains were administered via oral gavage and the next day mice were scanned with ultrasound using the setup depicted in **Fig S8a**. One day later on day 3, mice were sacrificed and their intestines were fixed for histology. (**d-e**) Colony forming units (CFU) per gram of feces (d) and total BURST* ultrasound signal imaged (e) on day 2 of piperacillin treatment (100 or 200 µg/mL) for mice colonized by thsS(t3)R-Bxb1_P7-bARG_Ser_ (bARG_Ser_) or thsS(t3)R-Bxb1_P7-GFP_mCherry (GFP) EcN. (**f**) Ultrasound images overlaying the integrated BURST* signal onto the integrated B-mode images for all mice on day 2 of piperacillin treatment. (**g**) Histopathology scoring of cecal tissues from piperacillin-treated mice on day 3 of treatment broken down by category of abnormality. Mice which received 200 ug/mL piperacillin did not exhibit significantly more signs of disease than mice which received 100 ug/mL piperacillin, but both piperacillin-treated groups exhibited more signs of disease than chloramphenicol-treated mice (see **Fig. S15a**). (**h**) Representative images of H&E-stained sections of cecal tissue on day 3 of piperacillin treatment. Abnormalities are indicated in red: mucosal epithelial cell death and degeneration (box), mucosal crypt hyperplasia (circle), mucosal/submucosal edema (asterisk), mononuclear infiltrates (arrow), granulocytic infiltrates (arrowhead), dilated lymphatic (star).

**Figure S19:**
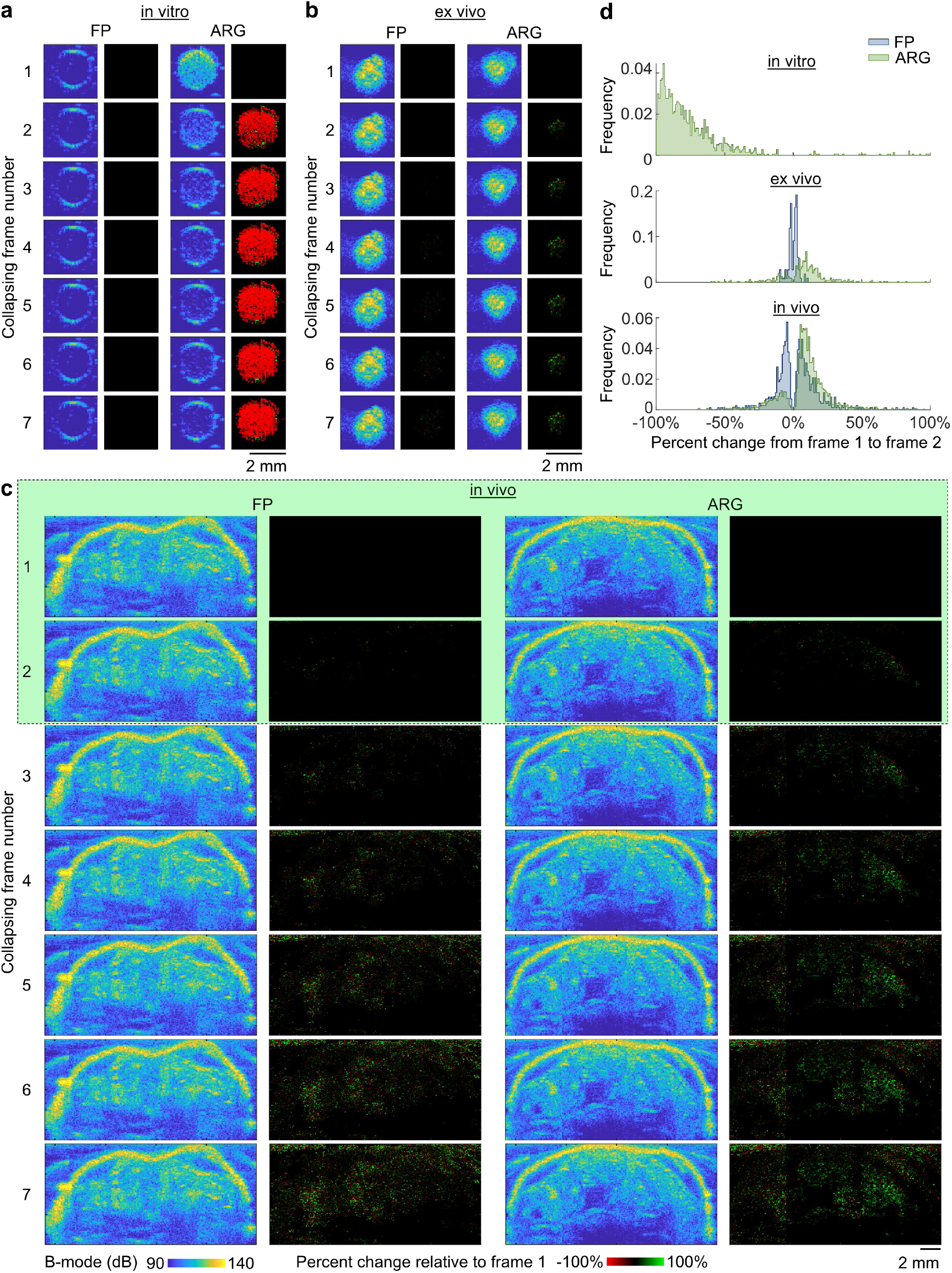
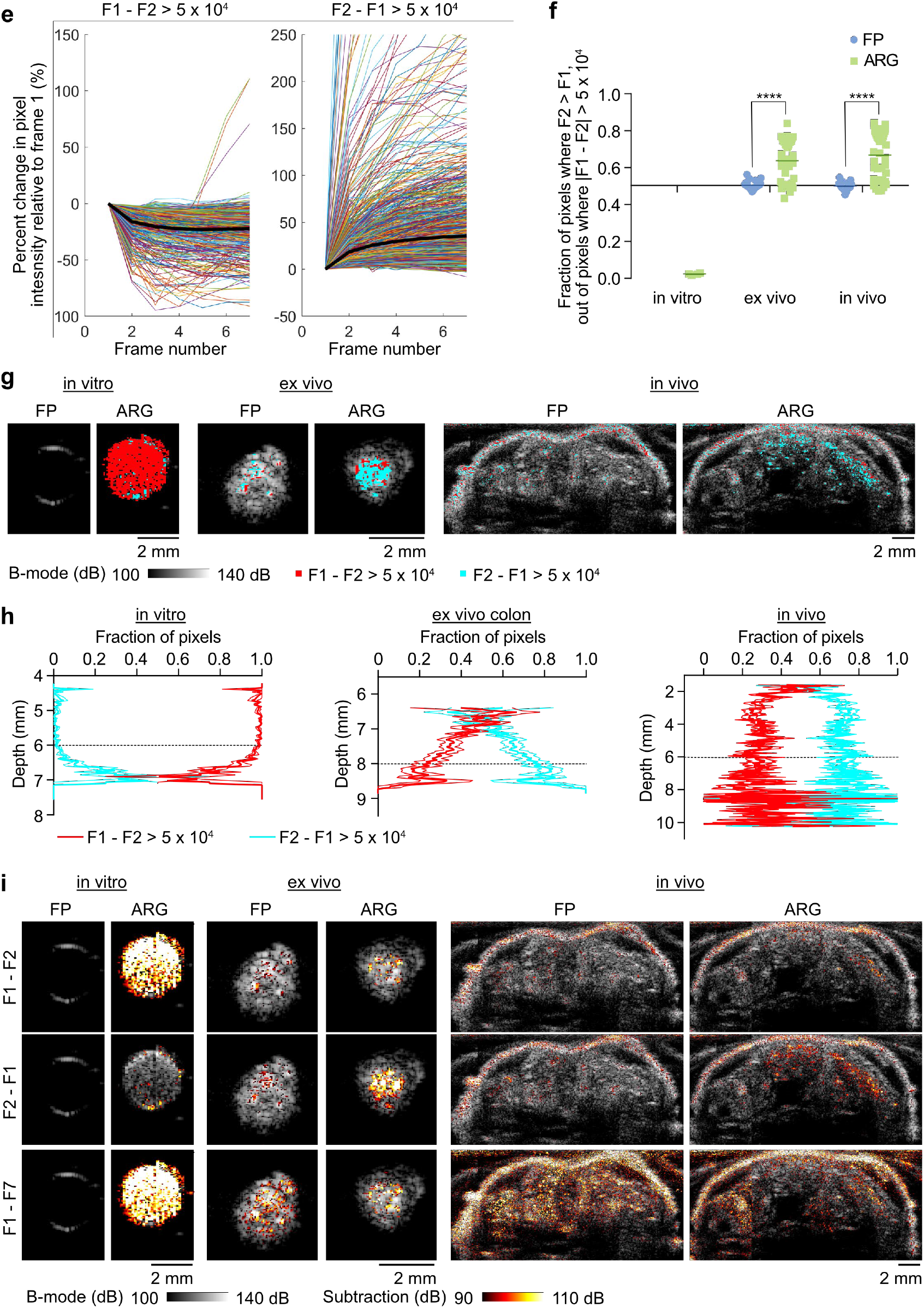
Comparison of processing BURST images in vitro, ex vivo, and in vivo. (**a-c**) B-mode images of the 7 collapsing frames (left) and percent difference images relative to the first collapsing frame (right) acquired using a rapid BURST script (that uses 3 focused beams at a time) of representative in vitro samples of EcN expressing bARG_Ser_ or a fluorescent protein (FP) at 5 × 10^8^ cells/mL (a), ex vivo colons colonized by bARG_Ser_ or FP-expressing EcN (b), and in vivo intestines of mice colonized by bARG_Ser_ or FP-expressing EcN (c). Color bars are the same for (a-c). To exclude noise, for the percent difference images, only pixels that had an absolute difference of 5 × 10^4^ or greater relative to the first collapsing frame were included. For (c), only the difference between the first two collapsing frames (green box) is useful due to tissue motion. (**d**) Histograms of the percent changes in pixel intensity from the first collapsing frame (frame 1) to the second collapsing frame (frame 2) for the representative in vitro, ex vivo, and in vivo images from (a-c). Only pixels that had an absolute difference of 5 × 10^4^ or greater were included; the FP in vitro images did not have any of these pixels. (**e**) Example pixel traces in terms of percent difference relative to frame 1 over the seven collapsing frames for the representative in vivo ARG acquisitions, categorized into pixels where the signal in the first collapsing frame (F1) was greater than the second (F2) by more than 5 × 10^4^ (F1 - F2 > 5 × 10^4^, left), and the pixels where the signal from the second collapsing frame (F2) was greater than the first (F1) by more than 5 × 10^4^ (F2 - F1 > 5 × 10^4^, right). Thin colored lines represent individual traces and bold black lines represent the mean. (**f**) Fraction of pixels where F2 > F1 out of all pixels that had an absolute difference of greater than 5 × 10^4^ between frames 1 and 2 for representative in vivo, ex vivo, and in vivo samples containing FP- or bARG_Ser_-expressing EcN. For in vitro samples, each point represents a biological replicate (N=4); FP in vitro acquisitions did not have any pixels that had an absolute difference of greater than 5 × 10^4^ between frames 1 and 2 so no data points are shown for this category. For ex vivo and in vivo samples, each point represents a BURST acquisition at a different location in the same mouse/intestines where the number of pixels that had an absolute difference of greater than 5 × 10^4^ between frames 1 and 2 was greater than 500 (from left to right, N = 28, 42, 15, 41). Lines represent the mean. Asterisks represent statistical significance by unpaired Student’s t-tests (**** = p < 0.00001; p-values from left to right: 2.163525e-008, 4.295376e-006). (**g**) Representative overlay images of the pixels where F1-F2 > 5 × 10^4^ (red), and the pixels where F2-F1 > 5 × 10^4^ (cyan). (**h**) Fraction of pixels where F1-F2 > 5 × 10^4^ or F2-F1 > 5 × 10^4^ versus the depth from the transducer for representative in vitro samples, ex vivo colons, and in vivo intestines containing ARG-expressing EcN. Bold colored lines represent the mean and thin colored lines represent the standard error of the mean (N = 4 biological replicates for in vitro samples, N = 30 cross-sectional images of an ex vivo colon which each contained more than 200 pixels where |F1-F2| > 5 × 10^4^, and N = 35 cross-sectional images of intestines in vivo which each contained more than 500 pixels where |F1-F2| > 5 × 10^4^). Dashed black lines represent the transducer focus. (**i**) Representative overlay images of different subtraction BURST images (hot scale) onto the B-mode image (greyscale): the first collapsing frame minus the second (F1-F2), the second collapsing frame minus the first (F2-F1), and the first collapsing frame minus the last (F1-F7). The negative portion of the subtraction images were removed for conversion to dB, and then the subtraction images were thresholded at 90 dB for the overlay.

**Supplementary note 1**: **BURST* imaging**.

Ex vivo and in vivo imaging in our study was performed using BURST* imaging. A modification of conventional BURST was required to obtain robust signals specific to ARG expression in samples with high background due to GI tissue and luminal contents. For in vitro samples containing ARGs, the collapse of the gas vesicles was apparent from raw B-mode images acquired at the collapsing pressure of BURST (**Fig. S19a**). In contrast, for ex vivo and in vivo samples containing ARGs, the difference in pixel intensity over the collapsing frames was small relative to the background and was confounded by tissue motion in later frames (**Fig. S19b-c**). For example, pixels were distributed around -70% difference from frame 1 to frame 2 for a representative in vitro sample with ARGs, while pixels were distributed around -16% and +18% difference from frame 1 to frame 2 for a representative in vivo sample with ARGs (**Fig. S19d**). Furthermore, the pixel intensities mostly decreased for ARG-containing in vitro samples while the pixel intensities tended to increase for ARG-containing ex vivo and in vivo samples over successive collapsing frames (**Fig. S19d-e**). For pixels that displayed an absolute difference of 5 × 10^4^ or greater, on average 98% of these pixels decreased from frame 1 to frame 2 for in vitro samples containing ARGs, while on average 64% and 67% of these pixels increased from frame 1 to frame 2 for ex vivo and in vivo acquisitions containing ARGs (**Fig. S19f**). This asymmetry and magnitude of changes is absent in control ex vivo and in vivo acquisitions with cells expressing fluorescent proteins (FP) (**Fig. S19f**). Thus, the ARG-specific signal in the ex vivo and in vivo acquisitions was characterized by a higher fraction of pixels with an increase in intensity from frame 1 to frame 2 of more than 5 × 10^4^. This trend could at least partially be caused by shadowing, as the percentage of pixels where F2-F1 > 5 × 10^4^ tended to increase with the depth from the transducer (**Fig. S19g-h**), but further work is needed to investigate the mechanism behind this trend.

Based on these trends, we used the following subtraction-based methods to calculate BURST images. For in vitro samples, the last collapsing frame was subtracted from the first (F1-F7) because the first collapsing frame displayed the most signal while the last frame displayed the least signal, and there was no movement to confound the signal across the 7 frames (**Fig. S19i**). For ex vivo and in vivo acquisitions, the first collapsing frame was subtracted from the second (F2-F1) because the ARG-containing samples exhibited a high fraction of pixels with an increase in intensity from frame 1 to frame 2 whereas control samples did not (**Fig. S19i**), and analyzing temporally adjacent frames minimized tissue motion artifacts.

### Description of supplementary videos

Supplementary Video 1: BURST*/B-mode tomogram of an arabinose- and streptomycin-treated mouse colonized by pBAD-bARG_Ser_ EcN

Supplementary Video 2: BURST*/B-mode tomogram of an arabinose- and streptomycin-treated mouse colonized by pBAD-RFP EcN

Supplementary Video 3: BURST*/B-mode tomogram of a chloramphenicol-treated mouse colonized by thsS(t3)R-Bxb1_P7-bARG_Ser_ EcN

Supplementary Video 4: BURST*/B-mode tomogram of a streptomycin-treated mouse colonized by thsS(t3)R-Bxb1_P7-bARG_Ser_ EcN

Supplementary Video 5: BURST*/B-mode tomogram of a streptomycin-treated mouse colonized by thsS(t3)R-Bxb1_P7-GFP_mCherry EcN

## Notes

### Competing Interest Statement

The authors have declared no competing interest.

