## Supplementary figures and images for "Probiotic acoustic biosensors for noninvasive imaging of gut inflammation"

### Supplementary_Video_1.gif

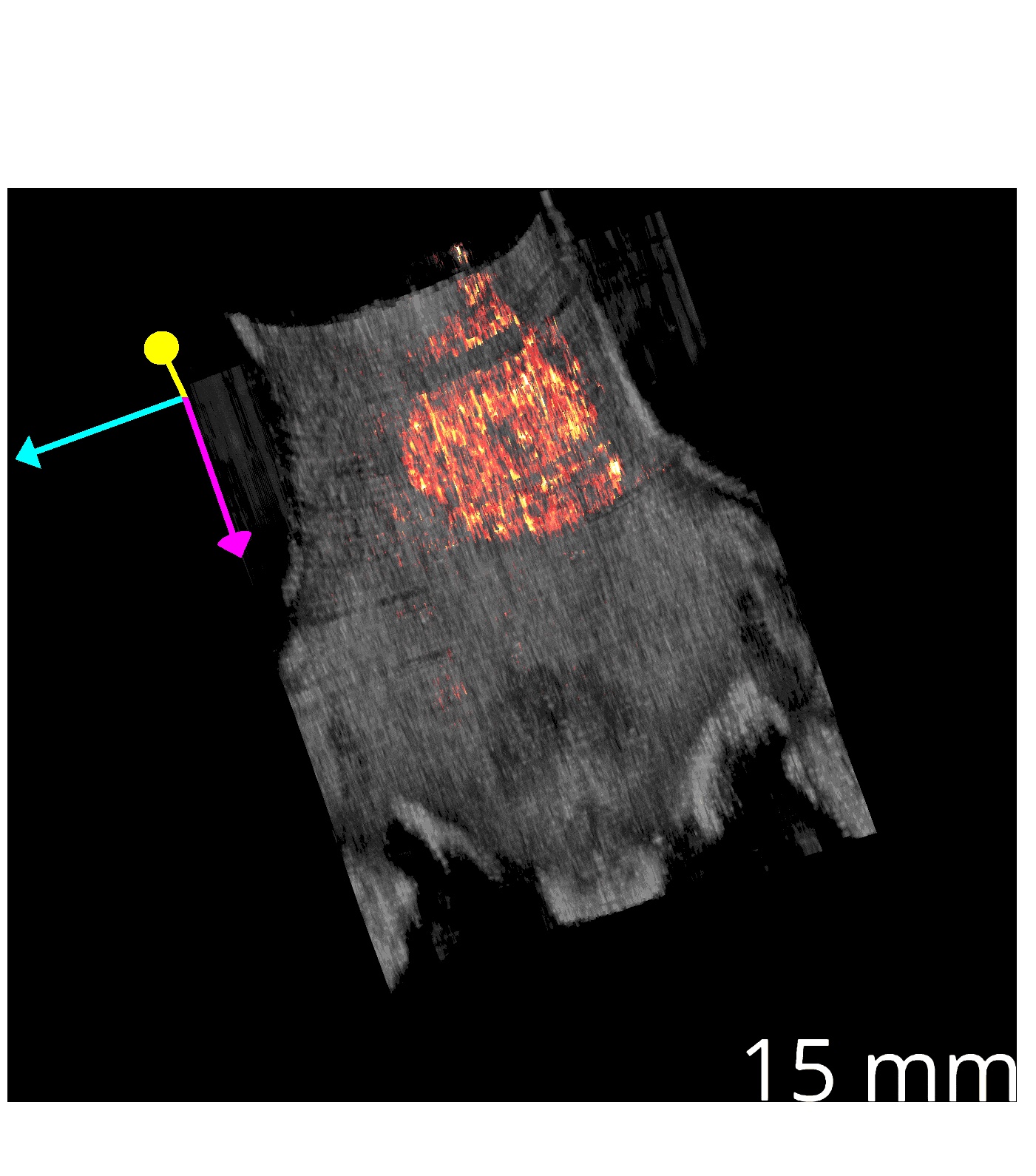

### Supplementary_Video_2.gif

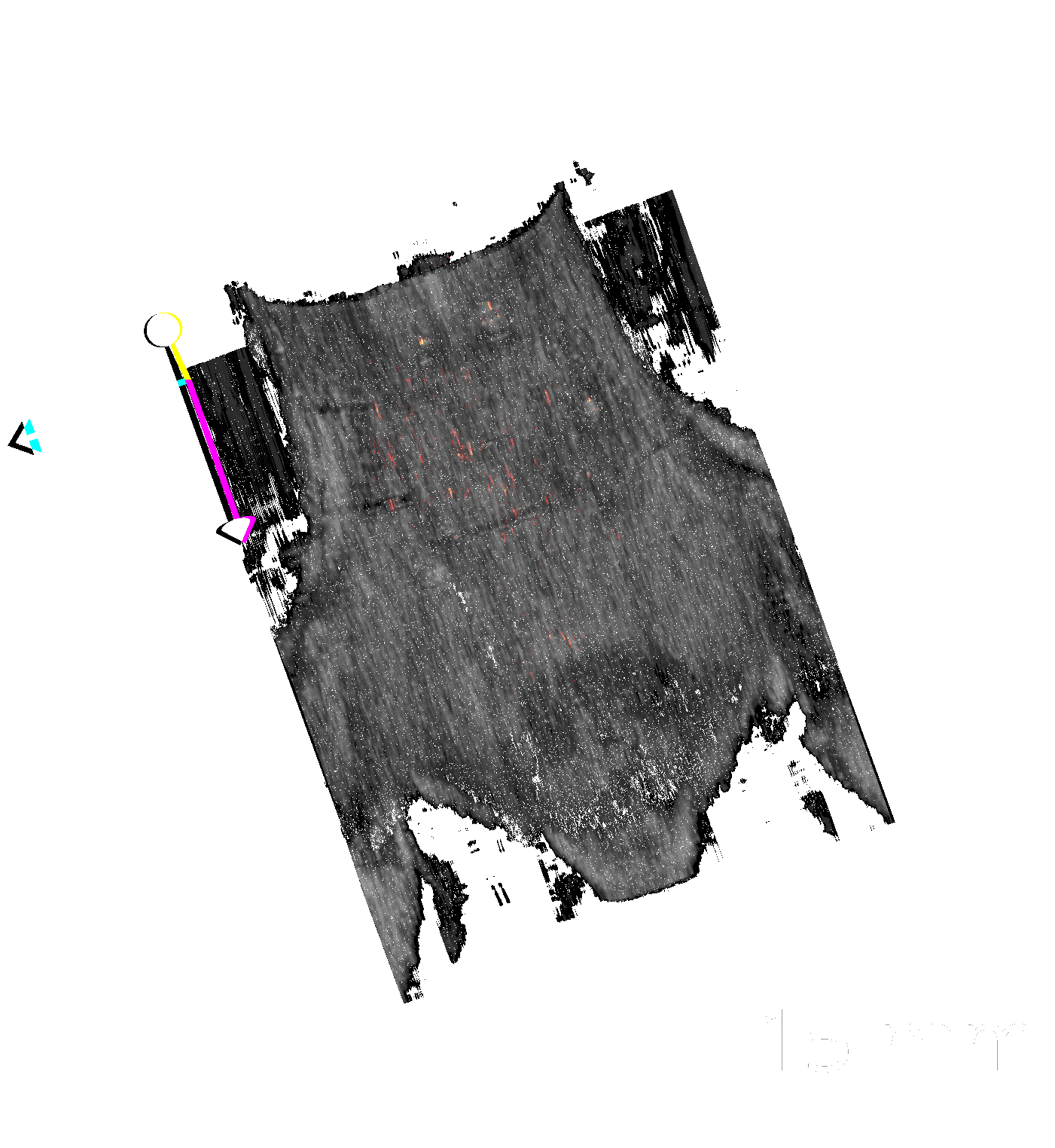

### Supplementary_Video_3.gif

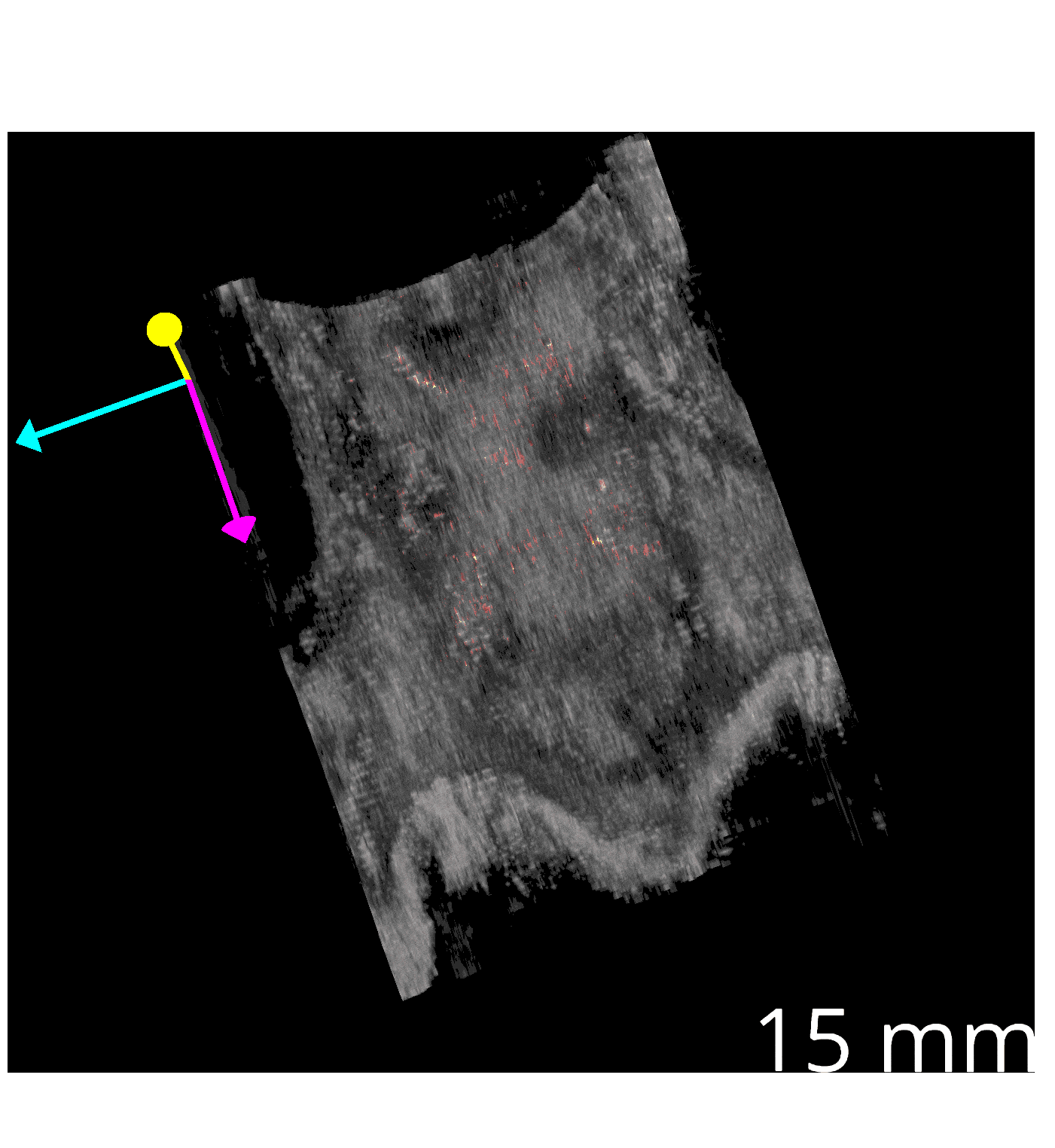

### Supplementary_Video_4.gif

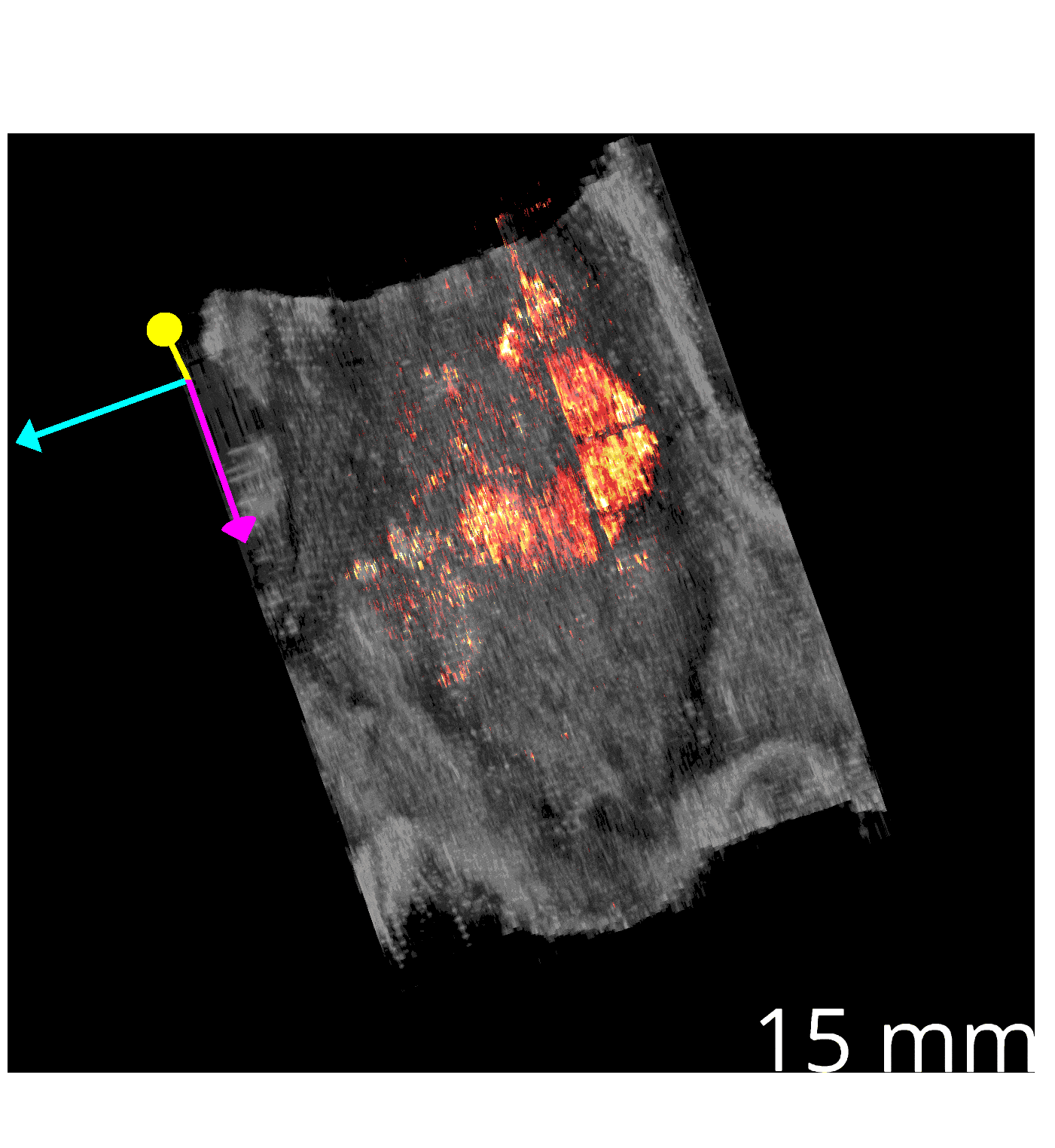

### Supplementary_Video_5.gif

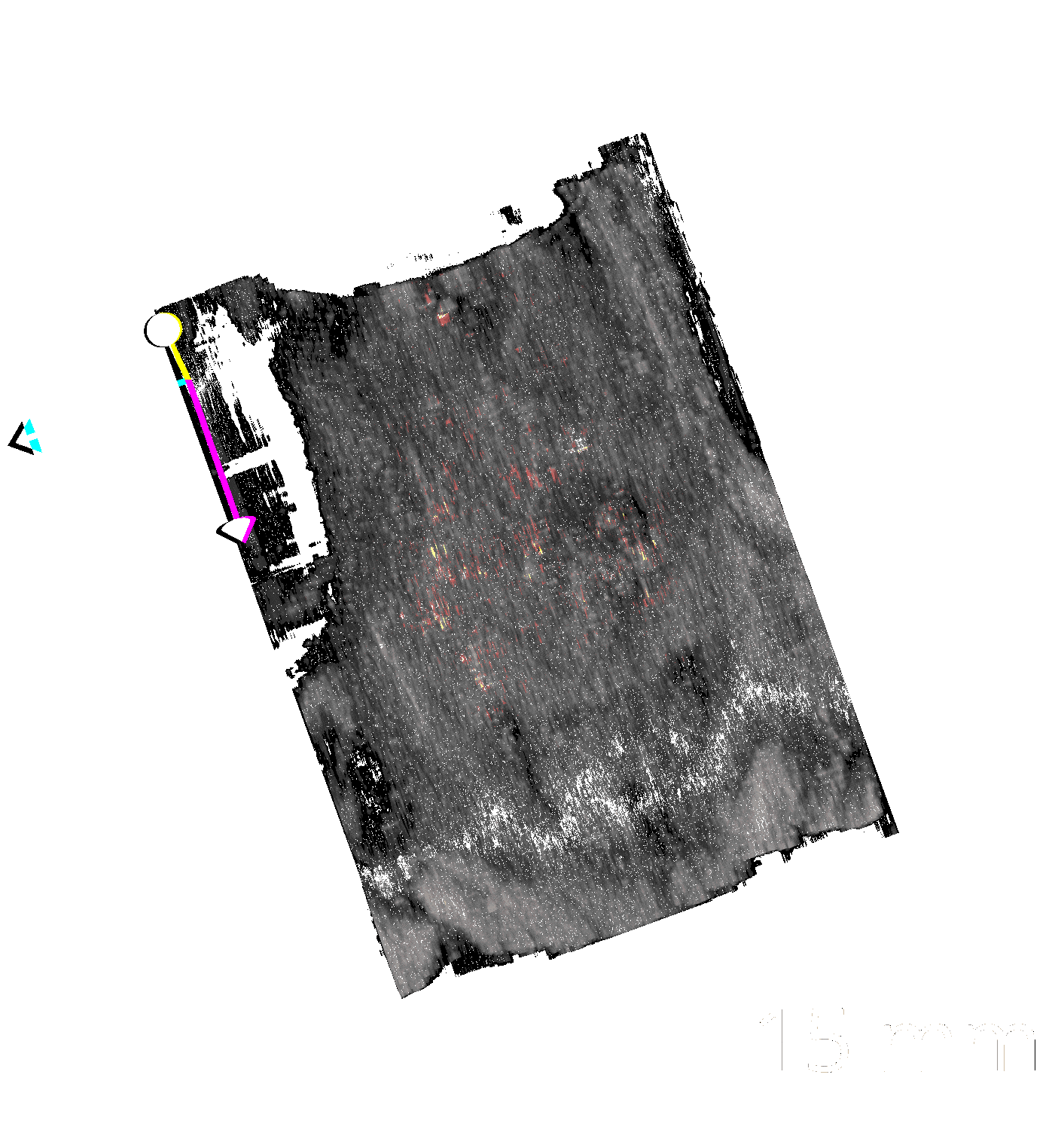
